# Disrupting *fzd9b* in zebrafish recapitulates stress- and anxiety-like behaviours associated with Williams syndrome

**DOI:** 10.1101/2024.01.22.576596

**Authors:** Jose V. Torres-Perez, Adele Leggieri, Aleksandra M. Mech, Sofia Anagianni, William Havelange, Caroline H. Brennan

## Abstract

Williams syndrome (WS) is a multifaceted developmental disorder characterized by a spectrum of physical and intellectual traits. Individuals with WS exhibit friendly, impulsive, and hyper-social behaviours, often coupled with anxiety. WS is attributed to a microdeletion on chromosome 7q11.23, affecting several genes, including *FZD9*, which plays an important role in neurodevelopment. Thus, we postulated that disruptions in *FZD9* might contribute to the behavioural features of WS including anxiety, and that pharmacological interventions targeting Wnt signalling, particularly the canonical pathway, might hold therapeutic potential for WS and related conditions. To test our hypothesis, we generated two mutant zebrafish lines with *fzd9b* disruptions. Our behavioural analysis revealed significant differences in stress- and anxiety-related responses at both larval and adult stages. Our attempt to restore stress reactivity by manipulating the Wnt/β-catenin pathway using a GSK-3 inhibitor was unsuccessful. Our qPCR data indicated a compensatory mechanism involving the upregulation of *fzd9b*, *wnt5b*, and *tafa5l* genes, potentially contributing to the observed phenotypes. These findings highlight the role of Fzd9b in modulating anxiety responses in zebrafish, offering potential avenues for novel therapeutics to address the neurological features of WS and related disorders.

**Summary statements:**

- We created mutant zebrafish lines to study stress reactivity and social behaviour, mirroring features found in Williams Syndrome (WS).
- Confirmation of *fzd9b* disruption revealed altered anxiety responses in larval and adult zebrafish.
- Marginal sociability increase was observed in the heterozygous fish for one of the two lines generated.
- Attempts to restore stress reactivity via the Wnt/β-catenin pathway manipulation were unsuccessful.
- We have identified compensation mechanisms involving upregulation of *wnt5b* and *tafa5l* genes.

## Introduction

Williams syndrome (WS), also known as Williams-Beuren syndrome, is a developmental disorder associated with a varying degree of intellectual disability and physical and behavioural features (Martens, Wilson and Reutens, 2008; Royston *et al*., 2021). Physical characteristics may include growth delay, cardiovascular disorders, distinctive facial traits, hypodontia and short stature. At a behavioural level, individuals with WS are described as friendly, empathic, highly impulsive, with short attention span, and hyper-social.

Despite the social predisposition, anxiety, especially that in the context of social scenarios, and anxiety-related disorders are a major comorbidity in WS (Ng-Cordell *et al*., 2018). Although rates vary enormously between studies, anxiety is considered the most prevalent mental health concern in WS (Royston *et al*., 2021). These altered anxiety and associated disorders are suggested to worsen with age (Stinton, Tomlinson and Estes, 2012; Ng-Cordell *et al*., 2018). Importantly, WS individuals’ responsiveness to anxiety situations seems related to modifications in the hypothalamic-pituitary-adrenal (HPA) axis, which affects their ability to control the intensity of their response, that is their reactivity, especially during socially loaded situations (Williams Syndrome Association, 2010; Lense and Dykens, 2013; Järvinen, Ng and Bellugi, 2015). Understanding WS anxiety biology can lead to interventions enhancing their quality of life.

WS is caused by a hemizygous microdeletion on chromosome 7q11.23. Most cases of WS occur as *de novo* mutations and affect males and females equally (Ewart *et al*., 1993; Morris, 1999). This deletion spans approximately 25-28 genes, including several genes with established developmental roles such as *BAZ1B*, *STX1A* and *FZD9* (Zhou *et al*., 2022), which makes it challenging to determine the role of each of those genes in relation to specific WS-associated features. Nonetheless, the study of individuals with atypical deletions, where one or more of the typically deleted genes persist, has unveiled genotype-phenotype correlations (Fusco et al., 2014; Chailangkarn et al., 2016). Fusco’s study revealed that individuals with intact genes near the proximal end of the classical deletion, including *FDZ9* and *BAZ1B* among others, exhibited milder craniofacial and cognitive characteristics thus suggesting their involvement in WS neurodevelopmental aspects (Fusco *et al*., 2014).

In a separate study, Chailangkarn and colleagues induced pluripotent stem cells (iPSCs) from neurotypical and WS individuals, including a WS subject that retained function for *FZD9 (*pWS88), to induce them into neural progenitor cells and cortical neurons (Chailangkarn *et al*., 2016). Focusing on neurons carrying markers for cortical layers V/VI associated with disrupted social behaviour (CTIP2-positive neurons), they found iPSC-derived CTIP2-positive neurons from WS individuals with the classical deletion had significantly higher total dendritic length, dendrite number and number of dendritic spines than iPSCs from typically developing individuals. Similar increased total cortical neuron dendritic length, dendrite number and spines are seen in postmortem brain of WS patients. Interestingly, CTIP2-positive neurons derived from the WS case with *FZD9* spared presented similar characteristics to neurons derived from typically developing subjects; thus, suggesting that lack of *FZD9* is responsible for the aberrant morpho-physiological characteristics observed in neuronal progenitors and mature neurons, of the classical WS cases.

*FZD9* encodes the transmembrane protein receptor frizzled 9, a member of the frizzled family of Wnt receptors (Wang *et al*., 1999). Fzd9 binds to Wnt proteins and has been associated with dendritic spine formation (Ramírez *et al*., 2016) and axonal growth (Shah *et al*., 2009); and mice lacking functional *Fzd9* show deficits in learning and memory tasks (Zhao and Pleasure, 2005). Expression of Fzd9 is first detected in neural precursor cells during the development of the neural tube, and remains high in the brain, heart, testis and skeletal muscle throughout development and into adulthood (Wang *et al*., 1999; Van Raay *et al*., 2001; Zhao and Pleasure, 2005).

In addition to an importance in neurodevelopment, disrupted Wnt signalling is implicated in different psychiatric and neurological disorders including bipolar disorders and schizophrenia, for which anxiety alterations have a high prevalence (Okerlund and Cheyette, 2011; Noelanders and Vleminckx, 2017). Current evidence also links Wnt signalling pathway dysregulations with altered stress responses associated with depression-like behaviours (Zhou *et al*., 2016) and attention-deficit hyperactivity disorders (Yde Ohki *et al*., 2020). Wnt signalling has been shown to modulate the response to stressors both at the level of beta catenin expression within the nucleus accumbens and the hippocampus, and behaviourally (Korem *et al*., 2017; Zwamborn *et al*., 2018). Furthermore, behavioural characterization showed that suppression of GSK3b (a key player in the canonical Wnt signalling pathway) in these brain areas has anxiolytic and pro-social effects (Latapy *et al*., 2012). Potentially relevant for therapy, Korem et al. demonstrated in a mouse shock and extinction stress model that the modulation of the canonical Wnt/β-catenin pathway with sulindac or LiCl influenced behavioural responses to the stressor and recovery from it (Korem *et al*., 2017).

Taking all together, we hypothesised that disruption of FZD9 signalling may underlie the anxiety-related phenotypes associated with WS and that pharmacological treatments aiming to restore Wnt signalling, specially targeting the canonical pathway, could be of therapeutic use for WS and related disorders. To test our hypothesis, we used zebrafish (*Danio rerio*), a small freshwater fish with a conserved neuro-endocrinal circuitry (de Abreu *et al*., 2021) and a high percentage of genetic homology (Bradford *et al*., 2017) to other vertebrates. Recent evidence indicates that deficits in *baz1b* may be more closely linked to the social aspects of WS (Torres-Pérez *et al*., 2023), thus we also hypothesized that *fzd9b* loss-of-function mutants might exhibit minimal changes in social behaviour. Although zebrafish have two paralogs for *fzd9* (*fzd9a* and *fzd9b*), only *fzd9b* is functionally relevant for the formation of the neural tube (Nikaido, Law and Kelsh, 2013). Therefore, we generated zebrafish with predicted loss-of-function mutations within *fzd9b* and assessed their social behaviour and anxiety-related responses compared to corresponding controls. As we observed differences in anxiety-related behaviours, we tested whether restoring the Wnt/β-catenin pathway with an acute treatment with 6-bromoindirubin-3’-oxime (BIO-acetoxime), a potent and selective GSK-3 inhibitor (Aourz *et al*., 2019), could revert the altered stress reactivity.

## Results

### CRISPR/Cas9-mediated disruption of *fzd9b* in zebrafish

Targeted mutations at the *fzd9b* gene were induced with the CRISPR/Cas9 system. Using this technique, two stable zebrafish lines were generated: a) *fzd9b^22bpDEL^*, a 22-base pair (bp) deletion that leads to a frameshift that, on translation, generates 4 amino acids (aa) followed by a premature stop codon (Fig. 1A-B); and b) *fzd9b^52bpDEL^*, a 52 bp deletion leading to a change in 44 aa also followed by a premature stop codon (Fig. 2A-B).

**Figure 1.**
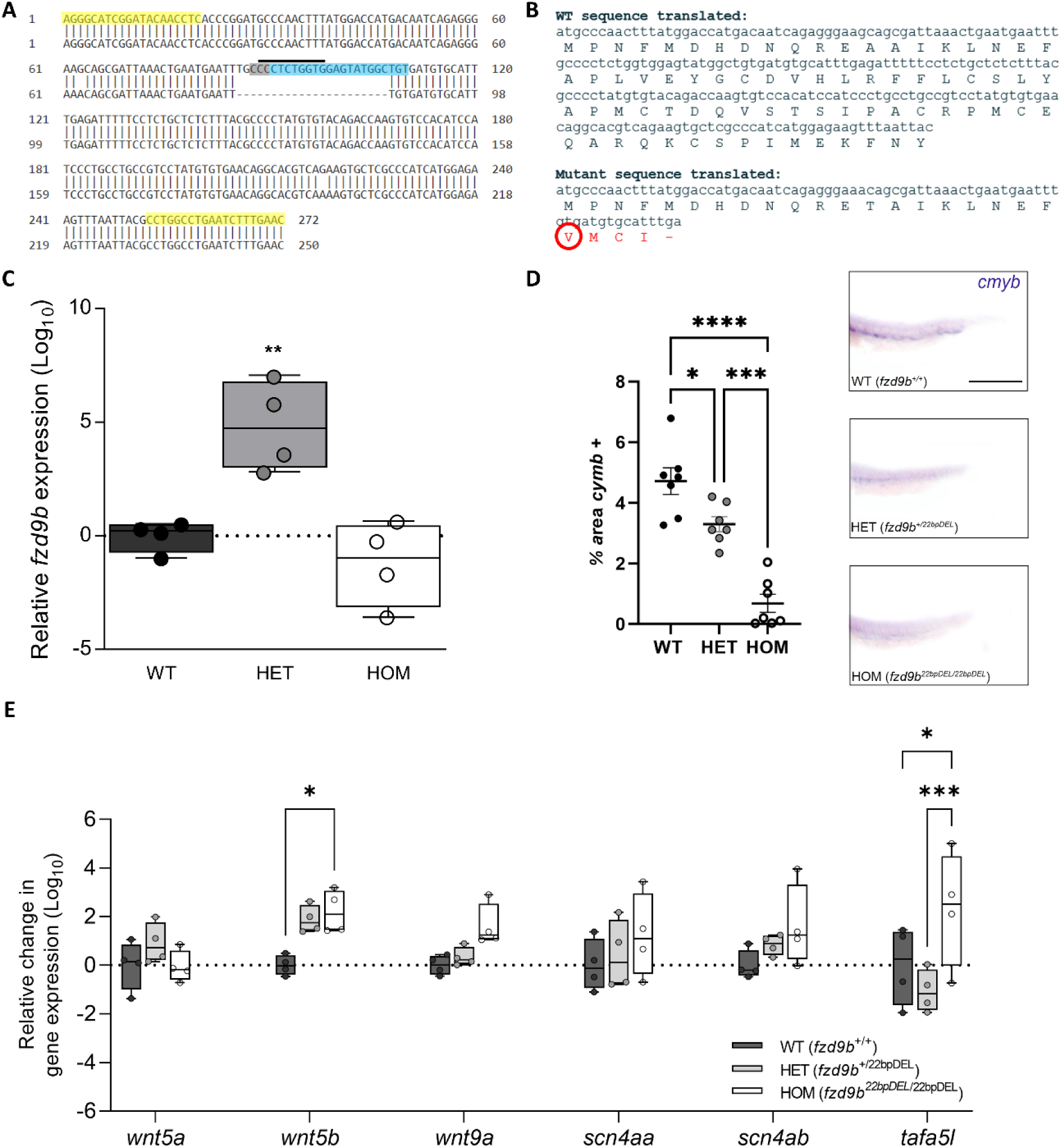
Characterization of the *fzd9b* 22bp deletion (*fzd9b^22bpDEL^*) line. **A)** Left panel shows a DNA blast for the portion of *fzd9b*’s exon 1 amplified by PCR used for genotyping between WT (above) and mutants (below, 22 bp deletion). Primers used are highlighted in yellow, crRNA site in blue, PAM site in grey and restriction enzyme over-lined. **B)** In-frame translation to amino acids (aa) sequences in WT (above) and mutants (below) of the same exon 1 portion. Red circle indicates mutation starting site, changed aa are in red and stop codon marked as a dash (-). **C)** Relative change in gene expression (log_10_) assessed by qPCR between 5 dpf larvae from each genotype showing that in HET the expression of *fzd9b* is significantly increased with respect to WT. Figure shows individual values (N = 4 per genotype, each representing a group of 16 larvae combined) and mean ± standard error mean (SEM). **D)** Quantification of the percentage of the whole body *in situ* hybridization (ISH) against *cmyb* at the tail of 40 hpf embryos from the three genotypes (N = 7); right panel shows representative embryos. **E)** qPCR analysis of the relative change in gene expression for *wnt5a*, *wnt5b*, *wnt9a*, *scn4aa*, *scn4ab* and *tafa5l* for pools of 5 dpf larvae (same as panel C) for the three genotypes. Graphs show box and whiskers with mean (central horizontal line) and minimum to maximum points. Dots represent biological replicates. In all cases: * *p* < 0.05, ** *p* < 0.01, *** *p* < 0.001, **** *p* < 0.000 vs. corresponding pairwise comparison. Scale bars represent 25 µm.

**Figure 2.**
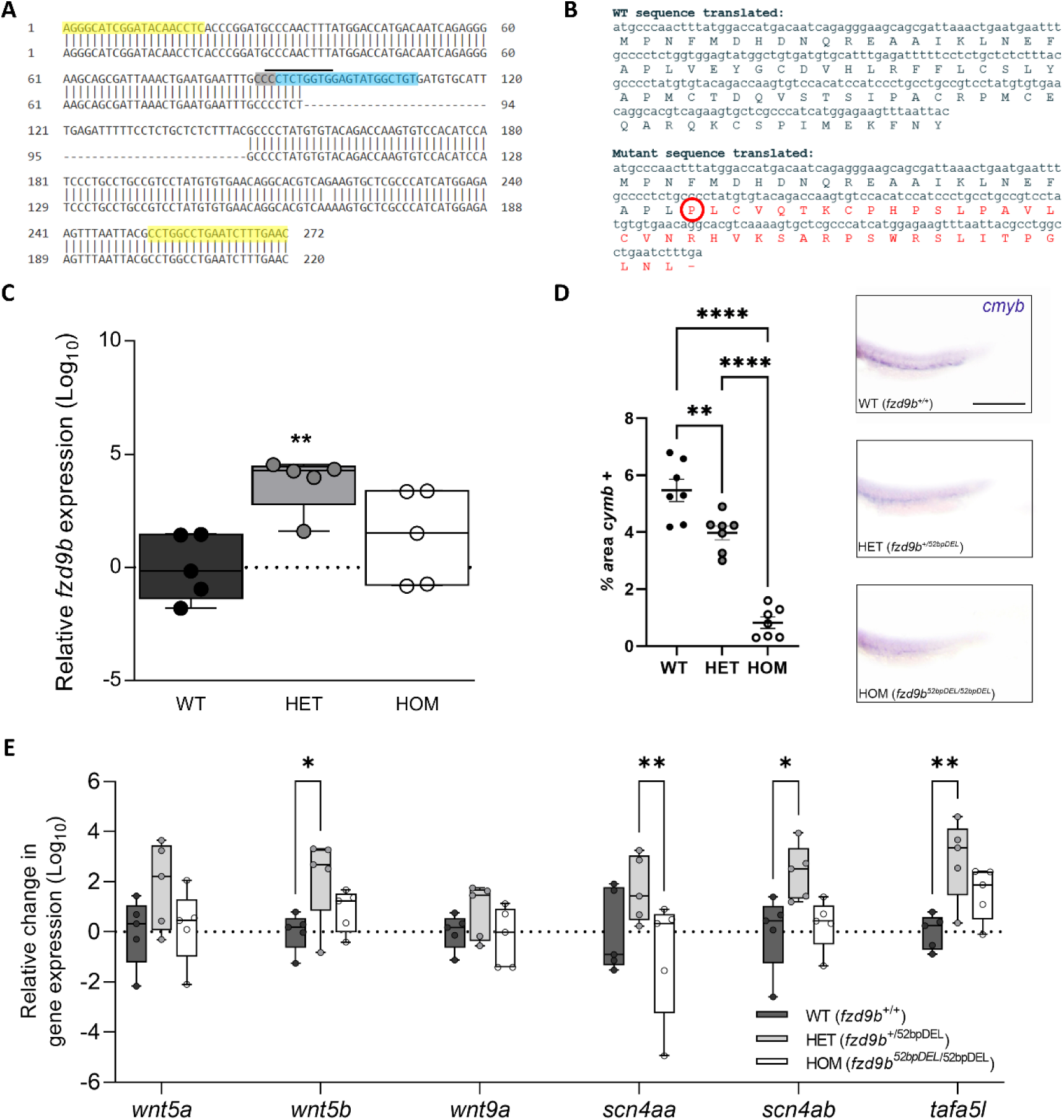
Characterization of the *fzd9b* 52bp deletion (*fzd9b^52bpDEL^*) line. **A)** Left panel shows a DNA blast for the portion of *fzd9b*’s exon 1 amplified by PCR used for genotyping between WT (above) and mutants (below, 52 bp deletion). Primers used are highlighted in yellow, crRNA site in blue, PAM site in grey and restriction enzyme over-lined. **B)** In-frame translation to amino acids (aa) sequences in WT (above) and mutants (below) of the same exon 1 portion. Red circle indicates mutation starting site, changed aa are in red and stop codon marked as a dash (-). **C)** Relative change in gene expression (log_10_) assessed by qPCR between 5 dpf larvae from each genotype showing that in HET the expression of *fzd9b* is significantly increased with respect to WT. Figure shows individual values (N = 5 per genotype, each representing a group of 16 larvae combined) and mean ± standard error mean (SEM). **D)** Quantification of the percentage of ISH against *cmyb* at the tail of 40 hpf embryos from the three genotypes (N = 7); right panel shows representative embryos. **E)** qPCR analysis of the relative change in gene expression for *wnt5a*, *wnt5b*, *wnt9a*, *scn4aa*, *scn4ab* and *tafa5l* for pools of 5 dpf larvae (same as panel C) for the three genotypes. Graphs show box and whiskers with mean (central horizontal line) and minimum to maximum points. Dots represent biological replicates. In all cases: * *p* < 0.05, ** *p* < 0.01, *** *p* < 0.001, **** *p* < 0.0001, vs. corresponding pairwise comparison. Scale bars represent 25 µm.

To quantify the amount of *fzd9b* mRNA in both lines, we used quantitative PCR (qPCR) with primers targeting *fzd9b* at 5’, prior to the mutation site. In both cases, the ANOVA statistical assay showed a significant difference in the expression of this mRNA between genotypes (*fzd9b^22bpDEL^*: F (2, 9) = 15.95, p = 0.0011; *fzd9b^52bpDEL^*: F (2, 12) = 6.866, p = 0.0103, Fig 1C), and post-hoc analysis revealed it to be driven by a significant increase in the heterozygous (*fzd9b^+/-^*, HET) when compared to wildtype (*fzd9b^+/+^*, WT) siblings (HET vs. WT; *fzd9b^22bpDEL^*: p = 0.0021, Fig. 1C; *fzd9b^52bpDEL^*: p = 0.0033, Fig. 2C) and homozygous (*fzd9b^-/-^*, HOM; HET vs. HOM; *fzd9b^22bpDEL^*: p = 0.0005; *fzd9b^52bpDEL^*: p = 0.0370), with no other comparisons reaching significance (all p > 0.05). Since *fzd9b* is a single exon gene and the mRNA surveillance pathway requires aberrant splicing during mRNA maturation to target degradation (Nicholson and Mühlemann, 2010), this suggests that both complete Fzd9b and smaller truncated proteins (3D structure in Sup. Fig. 1) are being expressed.

The levels of functional Fzd9b during zebrafish development are known to affect the development of haematopoietic stem cells (HPSC) (Grainger *et al*., 2019). Accordingly, we used ISH against *cmyb*, a marker of HPSC expansion (Grainger *et al*., 2019), above the yolk sack extension of 40 hours post fertilization (hpf) embryos, to assess downstream effects of disrupting *fzd9b*. One-way ANOVA followed by Dunnett’s multiple comparisons test revealed a significant decrease in *cymb* expression in *fzd9b* mutants compared to WT siblings in both *fzd9b^22bpDEL^* (WT vs HET p = 0.0219; WT vs HOM p < 0.0001, HET vs HOM p_adj_ = 0.0001; Fig 1D) and *fzd9b^52bpDEL^* (WT vs HET p = 0.0050; WT vs HOM p < 0.0001, HET vs HOM p < 0.0001; Fig 2D) lines.

### Disrupting *fzd9b* leads to changes in the expression of interacting genes

To further confirm the disruption of *fzd9b* in the zebrafish lines, we quantified the expression of genes suspected to interact with Fzd9b by qPCR: *wnt5a/b*, as WNT5A requires FZD9 to increase dendritic spine density in rodents (Ramírez *et al*., 2016); *wnt9a*, which is known to interact with Fzd9b in zebrafish (Grainger et al., 2019)(Grainger et al., 2019); and *scn4aa/b* (sodium channel, voltage gated, type IV alpha subunits) and *tafa5l* (syn. *FAM19A5*), the expression of which is affected in neural progenitor cells and neurons, respectively, derived from iPSCs from the classical WS patients but spared in the pWS88 individual, where *FZD9* is not missing (Chailangkarn *et al*., 2016).

In the *fzd9b^22bpDEL^* line, two-way ANOVA revealed a significant main effect of ‘genotype’ (F (2, 9) = 4.386, P=0.0468) and significant effect of the interaction between ‘gene of interest’ and ‘genotype’ of the animals (F (10, 45) = 2.689, p = 0.0114). Post-hoc analysis with Bonferroni’s correction for multiple testing showed that *wnt5b*, but not *wnt5a*, was significantly upregulated in the HOMs when compared to WT siblings (p = 0.0241; Fig 1E). Although it did not reach significance, HETs also showed a tendency for increased *wnt5b* expression (p = 0.0663; Fig. 1E). Similarly, there appeared to be an increase in *wnt9a* expression in hom, but this did not reach significance (p= 0.15). Additionally, there was a significant increase in the expression of *tafa5l* in the HOM fish (vs. WT: p = 0.0159; vs. HET: p = 0.0003; Fig 1E). No other post-hoc comparisons were significant for this line (p > 0.05).

In the other line, *fzd9b^52bpDEL^*, two-way ANOVA described a main effect of genotype (F (2, 12) = 6.140, p = 0.0146), and a main effect of gene of interest (F (5, 60) = 3.114, p = 0.0145) but no interaction between factors (F (10, 60) = 1.155, p = 0.3386). Post-hoc analysis revealed that HET fish had a significant increase in *wnt5b* (p = 0.0396; Fig. 2E), *scn4ab* (p = 0.0233) and *tafa5l* (p = 0.0037) when compared with WTs, and in *scn4aa* (p = 0.0093) when compared with HOMs. None of the other comparisons was significantly different (all p > 0.05).

### Experiment 1: Disrupting *fzd9b* has a marginal effect on sociability in zebrafish at 3 weeks of age

Increased sociability is a common feature in WS and, additionally, it is suggested to interfere with the anxiety response in WS (Martens, Wilson and Reutens, 2008; Williams Syndrome Association, 2010; Lense and Dykens, 2013; Järvinen, Ng and Bellugi, 2015; Ng-Cordell *et al*., 2018; Royston *et al*., 2021). To assess if *fzd9b* participates in zebrafish social abilities, zebrafish were assessed in a ‘sociability assay’ at 3 weeks of age, when this behaviour is fully developed (Dreosti *et al*., 2015).

In both lines, regardless of genotype, Wilconox test showed that all genotypes had a preference to interact with either one (*fzd9b^22bpDEL^*: WT: -2368, HET: -2150, HOM: -2192; Fig. 3A; *fzd9b^52bpDEL^*: WT: -1931, HET: -2057, HOM: -2129; In all cases p < 0.001; Fig. 4A) or three (*fzd9b^22bpDEL^*: WT: -2246, HET: -2130, HOM: -2207; *fzd9b^52bpDEL^*: WT: - 1773, HET: -2105, HOM: -1837; In all cases p < 0.001) conspecifics.

**Figure 3.**
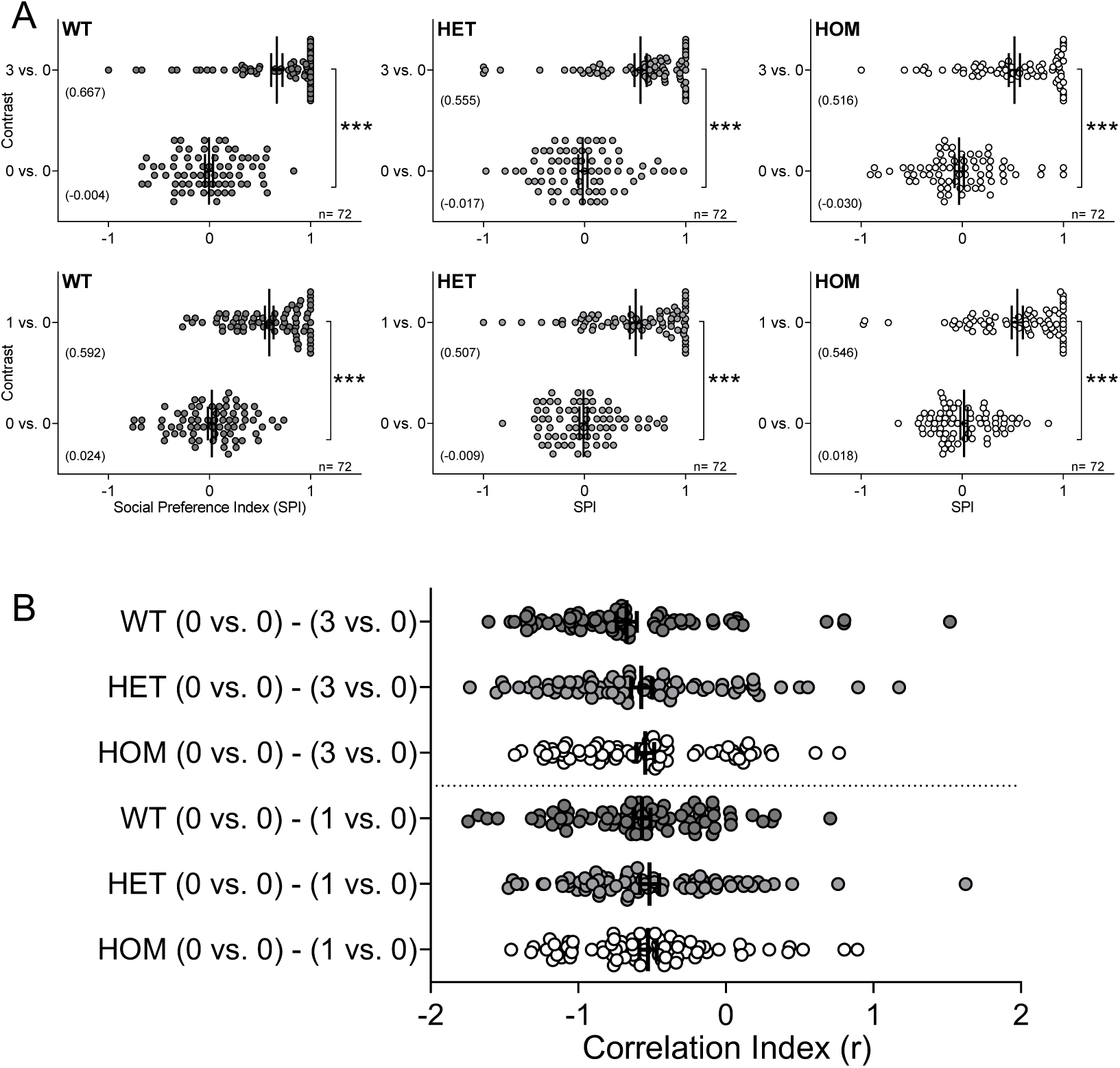
In the *fzd9b^22bpDEL^* line, sociability is not affected in 3-week old zebrafish. **A)** Graphs show change in social preference index (SPI) between basal conditions (0 vs. 0) and the 0 vs. 3 (first row) or 0 vs. 1 (second) contrast for either WT (*fzd9b^+/+^;* dark grey, first column), HET (*fzd9b^+/22bpDEL^*; light grey, second column) or HOM (*fzd9b^22bpDEL/22bpDEL^*; white, third column) at 3 weeks of age (between 20 to 22 dpf). Each graph includes corresponding N number and mean within brackets. **B)** Comparison of the correlation indexes (r) for all the contrast shown in A). Graphs depict individual values (circles of corresponding colour), mean ± SEM. In all cases: *** p < 0.001 vs. corresponding contrast.

**Figure 4.**
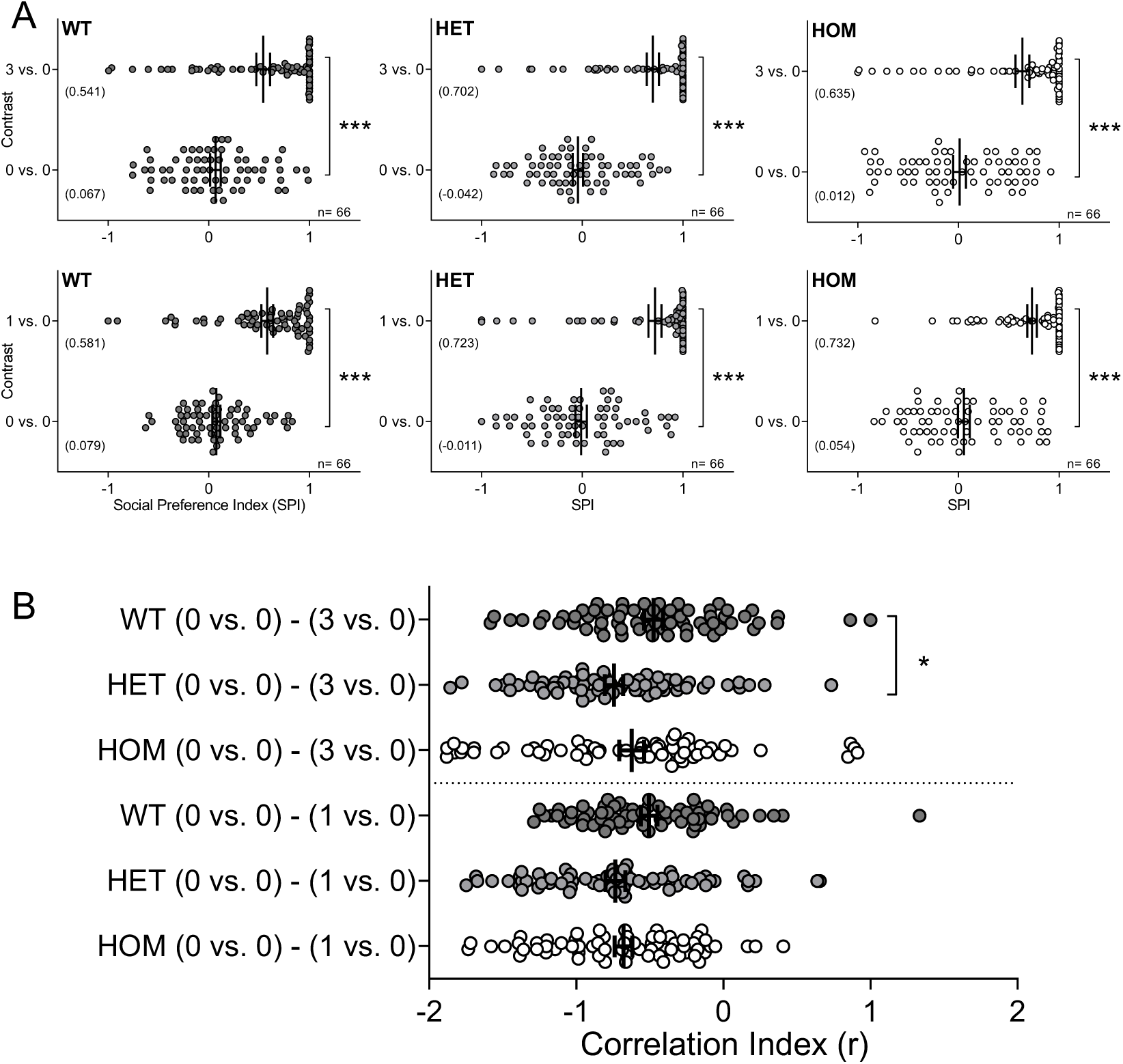
Sociability is marginally affected at 3-week of age in the zebrafish heterozygous from the *fzd9b^52bpDEL^* line. **A)** Graphs show change in SPI between basal (0 vs. 0) and the 0 vs. 3 (first row) or 0 vs. 1 (second) contrast for either WT (*fzd9b^+/+^;* dark grey, first column), HET (*fzd9b^+/52bpDEL^*; light grey, second column) or HOM (*fzd9b^52bpDEL/52bpDEL^*; white, third column) at 3 weeks of age (20-22 dpf). Graphs include n and mean. **B)** Comparison of ‘r’ for all the contrast shown in A). Graphs depict individual values (circles of corresponding colour), mean ± SEM. In all cases: *** p < 0.001 vs. corresponding contrast; * p < 0.05 vs. WT from the same contrast.

Additionally, to evaluate differences in their disposition to socialise between genotypes, we assessed the correlation indexes (r) (Torres-Pérez *et al*., 2023) by Kruskal-Wallis test. We found that there were no statistical differences by genotype in the *fzd9b^22bpDEL^* line (H (6) = 432, p = 0.2562; Fig. 3B). However, there was a significant difference in the *fzd9b^52bpDEL^* line (H (6) = 396, p = 0.0095; Fig. 4B) and post-hoc multiple comparisons with Dunn’s correction revealed this to be driven by HETs, showing a higher disposition to interact with multiple conspecifics (0 vs. 3), than WTs (p = 0.0476); with no other significant differences (all p > 0.05).

### Experiment 2: Lack of *fz9b* does not lead to changes in the response to forced changes in environmental light

Response to forced light dark transition (FLDT) was used to assess stress-related behaviour in zebrafish at 5 days post-fertilization (dpf). The forced light/dark transition paradigm (FLDT; Fig. 5), relies on the fish response to sudden change in environmental illumination and has previously been used to assess differences in stress-response (Peng *et al*., 2016; Lee *et al*., 2019; García-González *et al*., 2021; Faught and Vijayan, 2022)

**Figure 5.**
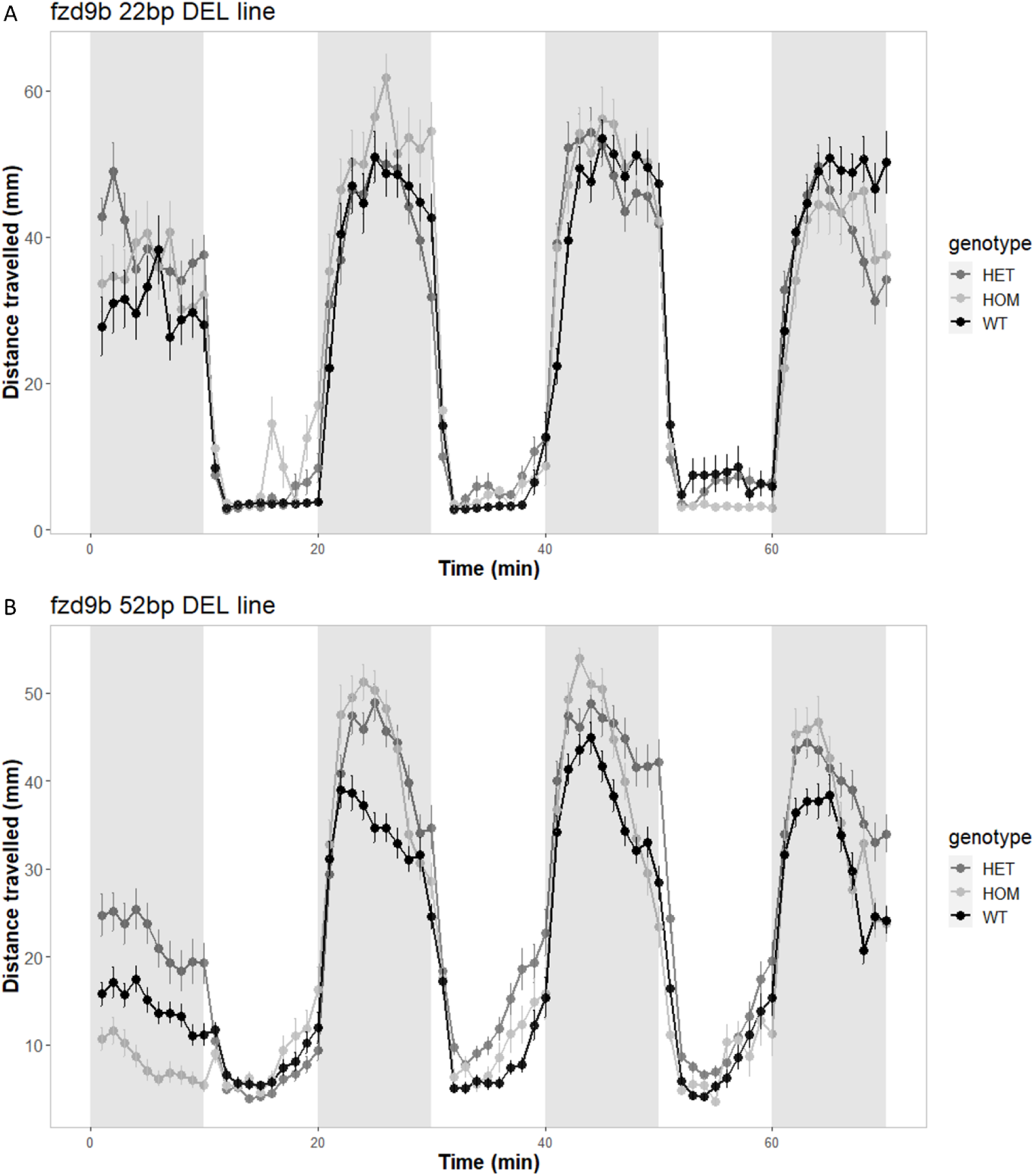
Disrupting *fzd9b* does not alter zebrafish responses in the forced light dark (FLD) assay. Mean distance moved per genotype during the baseline and the three light-dark cycles (10 minutes each) for **A)** the *fzd9b^22bpDEL^* line (n_WT_ = 48, n_HET_ = 93, N_HOM_ = 47) and, **B)** the *fzd9b^52bpDEL^* line (n_WT_ = 49, n_HET_ = 91, n_HOM_ = 48). Data shows mean ± SEM.

Results were similar for both lines (Fig. 5A for *fzd9b^22bpDEL^*; Fig. 5B for *fzd9b^52bpDEL^*). There was a main effect of time in baseline, light and dark periods with no effect of genotype and no genotype by time interaction (Supplementary Table 2 for p-values). In response to the transition from dark to light, fish display an initial period of freezing/reduced movement that gradually increases towards baseline levels over the 10 min period in the light. We assessed these changes in locomotion during the light period measured as the sum of slopes from the light periods. Again, there was no significant effect of genotype (Supplementary Table 2).

### Experiment 3: Lack of *fz9b* results in differential habituation in the repeated acoustic startle assay

Habituation to acoustic startle is used as a measure of associative learning (Basnet *et al*., 2019; Beppi, Straumann and Bögli, 2021; García-González *et al*., 2021) but magnitude of the response and degree of habituation can also be used as a measure of hyper-arousal associated with exaggerated stress responses (Cohen *et al*., 2006; Tanaka *et al*., 2019). We also included a brief flash of light at the start and end of the assay to determine whether potential differences in response were stimulus specific. We determined the baseline locomotion, the distance travelled in response to the flash of light before and after acoustic startle, and the maximum peak distance travelled in response to acoustic startle, mean peak distance across all taps, proportion of responders and rate of habituation in response to repeated acoustic startle using both distance travelled and proportion of responders for each stimulus for each genotype.

In the *fzd9b^22bpDEL^* line, as for FLDT, there was no significant effect of genotype on distance travelled prior to the first flash of light and prior to the repeated acoustic stimuli, with only time having a significant main effect (χ2 (8) = 24.0650, p < 0.05). During the tap events, we did not observe any effect of genotype on distance travelled, and only time had a significant main effect (χ2 (19) = 658.2177, p < 0.0001; Fig. 6A).

**Figure 6.**
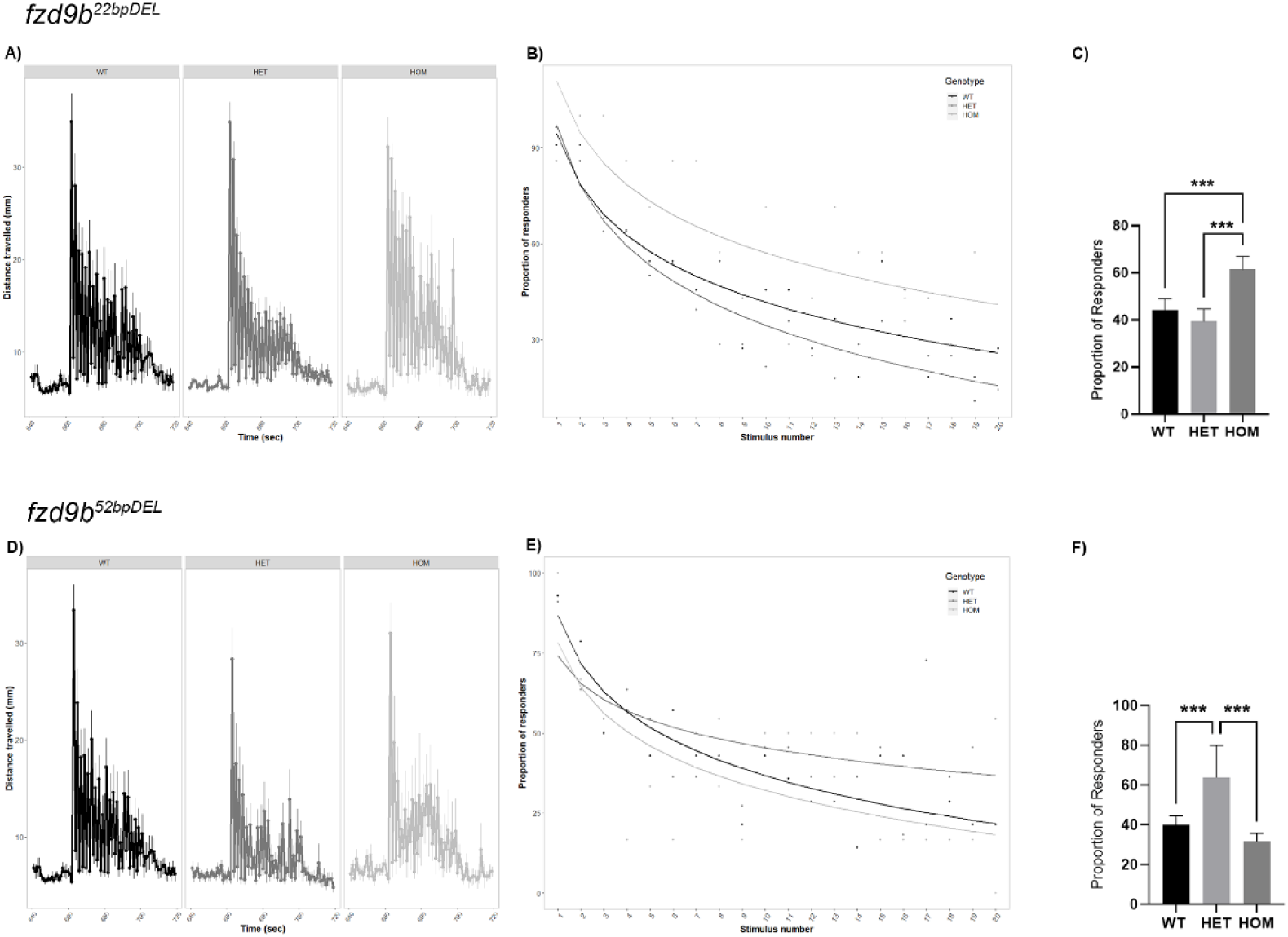
Disrupting *fzd9b* leads to slower habituation in the acoustic startle assay. A), B) and C) represent the *fzd9b^22bpDEL^* line (n_WT_ = 45, n_HET_ = 91, n_HOM_ = 43) and D), E) and F) represent the *fzd9b^52bpDEL^* line (n_WT_ = 46, n_HET_ = 85, n_HOM_ = 47). Response and habituation to acoustic startle of zebrafish larvae was assessed at 5 dpf. Paradigm: 10 min dark (baseline), 2 sec light, 1 min dark, 20 taps (2 sec interval in between). A) D) Mean distance moved during the acoustic cues. B) E) Bar plot for proportion of proportion of responders during the acoustic cues. There was a significant difference in the number of responders in B) showing that HOMs take longer to habituate and in E) that HETs take longer to habituate. C) F) Bar plot for the proportion of responders over all acoustic stimuli. Data shows mean ± SEM. Dots represent biological replicates. In all cases: * *p* < 0.05, ** *p* < 0.01, *** *p* < 0.001 vs. corresponding WT.

When assessing the proportion of responders, larvae showed a habituation response to repeated acoustic startle consistent with previous reports (García-González *et al*., 2021; Leggieri *et al*., 2022): 91% of WT zebrafish responded to the first acoustic stimulus, but only 25% of WT responded to the last. There was a significant main effect of genotype on proportion of responders (χ2 (2) = 56.126, p < 0.0001), and a significant two-way interaction between the stimulus events and the genotype (χ2 (22) = 1.411411, p < 0.0001), such that HOMs habituated significantly more slowly than WTs (p < 0.0001) and HETs (p < 0.0001; Fig. 6B-C).

During the flash of light stimuli, before and after the tap events, there were no significant differences between genotypes in neither the magnitude of response nor the slope of recovery (p > 0.05).

In a similar fashion, the *fzd9b^52bpDEL^* line, only time had a significant main effect on distance travelled prior to the first flash of light and the repeated acoustic stimuli (χ2(8) = 25.03250, p < 0.01) and during tap events (χ2 (19) = 192.7586, p < 0.0001; Fig. 6D); with genotype not having a significant main effect. Additionally, there were no significant differences between genotypes in neither the magnitude of response nor the slope of recovery (p > 0.05) during the flashes of light before and after the tap events.

When assessing the proportion of responders, the *fzd9b^52bpDEL^*line showed a habituation response to repeated acoustic startle similar to that reported in the previous line, ranging from 93% of WT zebrafish in response to the first acoustic stimulus to 21% of WT responding to the last. There was a significant main effect of genotype on proportion of responders (χ2 (2) = 11.212, p < 0.01) and a significant two-way interaction between the stimulus events and the genotype (χ2 (22) = 1.94786, p < 0.0001), whereby HET fish habituate more slowly than WTS (p < 0.0001) and HOMs (p < 0.0001) while there was no difference between WTs and HOMs (p > 0.05; Fig. 6E-F).

### Experiment 4: Adult fish mutant for *fzd9b* show altered anxiety-like behaviour as assessed by the novel tank diving assay

Anxiety is a common feature in WS which might be exacerbated by age (Stinton, Tomlinson and Estes, 2012; Ng-Cordell *et al*., 2018; Royston *et al*., 2021). Therefore, to assess if disruption of *fzd9b* affected this behaviour in adult zebrafish, we used the novel tank diving assay. In this assay, the time spent in, and distance to, the bottom of the tank is used to determine the level of anxiety-like phenotypes in zebrafish (Evans *et al*., 2021). In the *fzd9b^22bpDEL^* line, the analysis of the total distance travelled showed a significant effect of time (F (10) = 26.4103, p = 0.003) by which the first-time bin showed significantly lower locomotion than subsequent ones (1^st^ time bin vs. 4^th^-7^th^, 9^th^, 11^th^: p < 0.05; Fig. 6A). No significant effect of the interaction time-genotype (F (20) = 15.7835, p = 0.730) nor the factor genotype alone (F (2) = 3.4881, p = 0.175) was seen.

When analysing the distance from the bottom (Fig. 6B) there was a significant main effect of genotype (F (2) = 27.103, p < 0.001) driven by WTs having a significantly greater distance from the bottom than either HET or HOM overall (vs. HET: p < 0.001; vs. HOM: p < 0.001) and no main effect of time (F (10) = 15.923, p = 0.1019). There was a significant time x genotype interaction (F (20) = 97.432, p < 0.001) with post-hoc analysis indicating that WTs significantly reduced their distance from the bottom over time (1^st^ vs. 9^th^ time bin: p = 0.05; 2^nd^ vs. 9^th^: p = 0.018; vs. 10^th^: p = 0.033) while HETs increase their distance (1^st^ vs. 4^th^: p 0.002; vs. 5^th^-11^th^: p < 0.001) and HOMs did not show any significant trend.

The analysis of time spent on the bottom third (Fig 6C) showed that there was a main effect of genotype (F (2) = 213.4849, p < 0.001) where all genotypes differed from each other (all comparisons: p < 0.001) such that WTs spent significantly less time in bottom third than the other two genotypes, HOMs spent significantly more time in the bottom third than the other two genotypes, and HETs’ time in bottom third was somewhere in-between WTs and HOMs but differed significantly from both. There was no main effect of time (F (10) = 4.5732, p = 0.9178). The interaction time x genotype was also significant (F (20) = 39.2124, p < 0.001). Post-hoc analysis showed that, while there were no significant differences between time bins for WTs and HOMs, HETs showed a significant tendency to spend less time in the bottom third of the tank the longer they are in the novel environment (1^st^ time bin vs. 5^th^-7^th^: p < 0.05; vs. 9^th^-11^th^: p < 0.04). Finally, we also assessed the top half visits (Fig. 6D) where there was a significant effect of time (F (10) = 39.1785, p < 0.001), which showed that fish made more top half visits over time (1^st^ time bin vs. 4^th^-11^th^: p < 0.05), an indicator of reduced anxiety-like phenotype with time spent in the assay. Neither the factor genotype nor the interaction time x genotype were significant for top half visits (F (2) = 0.9664, p = 0.6168; F (20) = 21.845, p = 0.34898; respectively).

Similar analysis was carried out for the *fzd9b^52bpDEL^*line. For the total distance moved (Fig. 7A), there was a significant effect of time (F (10) = 279359, p = 0.002; driven by 2^nd^ time bin vs. 9^th^: p = 0.024) and the interaction time-genotype (F (20) = 34.3873, p = 0.024), while there was no main effect of genotype (F (2) = 1.9109, p = 0.3846). Post-hoc analysis showed that the time x genotype interaction was driven by HETs moving greater distances with time (1^st^ time bin vs. 8^th^-11^th^: p < 0.01) with no differences seen for WTs and HOMs. Similarly, the analysis of the distance from the bottom showed a significant main effect of time (F (10) = 87.1895, p < 0.001; Fig. 7B) where animals remain closer to the bottom in the initial time bins (1^st^ vs. 4^th^-11^th^: p < 0.001; 2^nd^ vs. 6^th^-11^th^: p < 0.05), but neither the genotype (F (2) = 5.7805, p = 0.056) nor the interaction (F (20) = 25.973, p = 0.167) were different.

**Figure 7.**
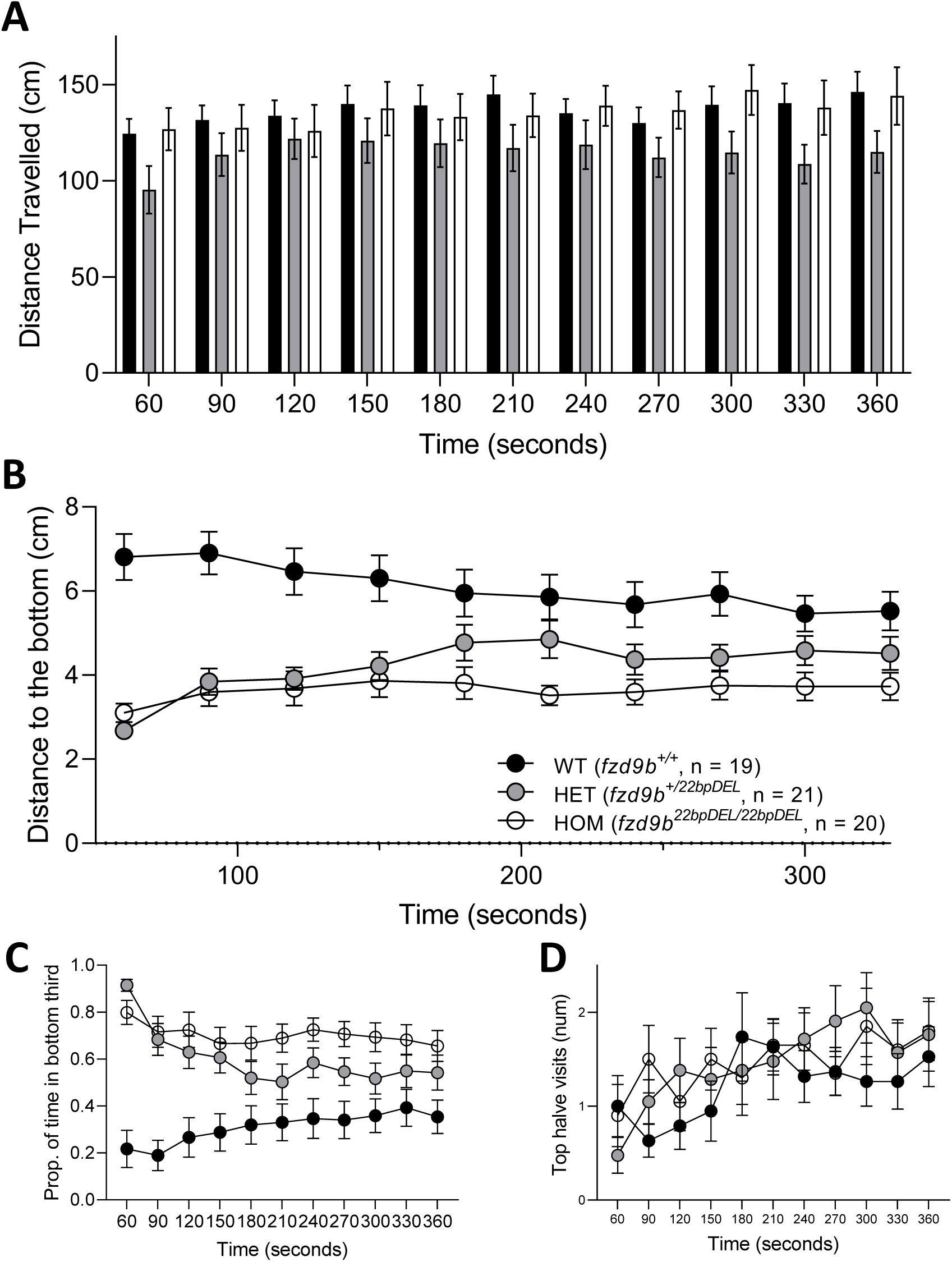
Novel tank diving assay shows reduced anxiety in the *fzd9b^22bpDEL^* line. **A)** Distance travelled for the three genotypes (WTs, HETs and HOMs) in 30 seconds time bins. **B)** Distance to the bottom for the three genotypes in similar time bins; graph includes n number. WTs showed a more anxious phenotype than both HETs and HOMs. **C)** Proportion of time in the bottom third, HETs had a tendency towards spending less time in the bottom third, and **D)** top halve visits. Graphs show mean ± SEM. Corresponding n numbers are shown in figure 6B.

The statistical analysis of the proportion of time spent on the bottom third for the *fzd9b^52bpDEL^* line (Fig. 7C) showed that there was a main effect time (F (10) = 32.550, p < 0.001), with reduced time in the bottom third through time (1^st^ time bin vs. 10^th^-11^th^: p < 0.05; 2^nd^ vs. 10^th^-11^th^: p < 0.05), and a significant main effect of genotype (F (2) = 29.611, p < 0.001), driven by the HETs spending less time on the bottom (vs. WT: p < 0.001; vs. HOM: p < 0.001; WT vs. HOM: p = 0.314), but not for the interaction time-genotype (F (20) = 14.567, p = 0.801). Finally, the analysis of top half visits in this zebrafish line (Fig. 7D) showed there was a main effect of time (F (10) = 25.8218, p = 0.004: driven by the 2^nd^ time bin being significantly different to the 10^th^ and 11^th^, p < 0.05) and the interaction time x genotype (F (20) = 45.6144, p < 0.001), while there was no statistical difference for the factor genotype (F (2) = 1.3812, p = 0.501). Post-hoc analysis showed that HETs and WT made more top half visits with time spent in the assay (HETS :1^st^ time bin vs. 4^th^-11^th^: p < 0.05, WT 1^st^ vs. 4^th^: p = 0.031), while there was no change in the HOMs.

### Experiment 5: Inhibition of GSK-3 does not restore stress-reactivity in zebrafish with disrupted *fzd9b*

To test our hypothesis that targeting the Wnt/β-catenin signalling canonical pathway could revert altered stress reactivity observed in the mutant *fzd9b* lines, we used an acute treatment with BIO-acetoxime, a selective GSK-3 inhibitor, at a dose of 200 µM, the maximal tolerated concentration that does not affecting locomotion in larval zebrafish (Aourz *et al*., 2019). GSK-3 has a prominent role within the Wnt/β-catenin signalling canonical pathway as part of the destruction complex that, under inactive conditions, degrades β-catenin and thus prevents its intracellular accumulation (Beurel, Grieco and Jope, 2015). Interestingly, inhibition of GSK3 might have a therapeutic potential in different neurological disorders and psychiatric diseases (Beurel, Grieco and Jope, 2015). As we saw no differences in larval FLDT assay across genotypes and cost precluded analysis of tank diving in adults, we used response to repeated acoustic startle as a measure of stress-reactivity.

After 90 minutes of exposure, zebrafish were subjected to an acoustic startle assay (Fig. 9A). Despite the potential impact of the drug on vision, we included, as with prior acoustic startle assay (Experiment 3), a brief flash of light at the start and end of the assay to determine whether potential differences in response were due to fatigue.

**Figure 8.**
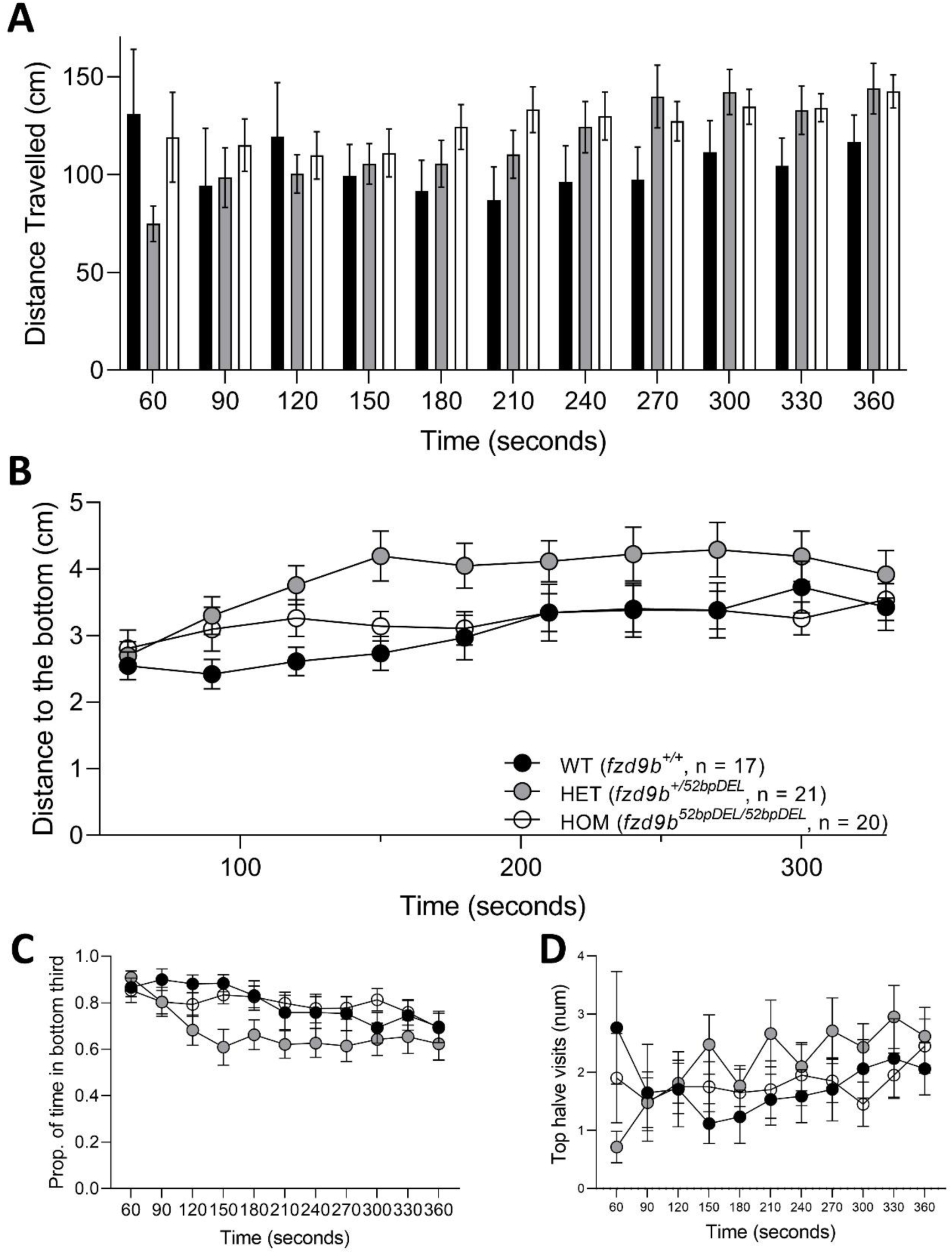
Novel tank diving assay shows mild differences in anxiety-like behaviours in the *fzd9b^52bpDEL^* line. **A)** Distance travelled, **B)** distance to the bottom, **C)** Proportion of time in the bottom third and **D)** top halve visits for the three genotypes (WTs, HETs and HOMs) in 30 seconds time bins. Although there was no difference by genotype in the distance to the bottom, the proportion of time in the bottom third for the HETs was significantly different than the other genotypes and HETs also visited significantly more the top half part of the tank. Graphs show mean ± SEM. Graph B includes the n number.

**Figure 9.**
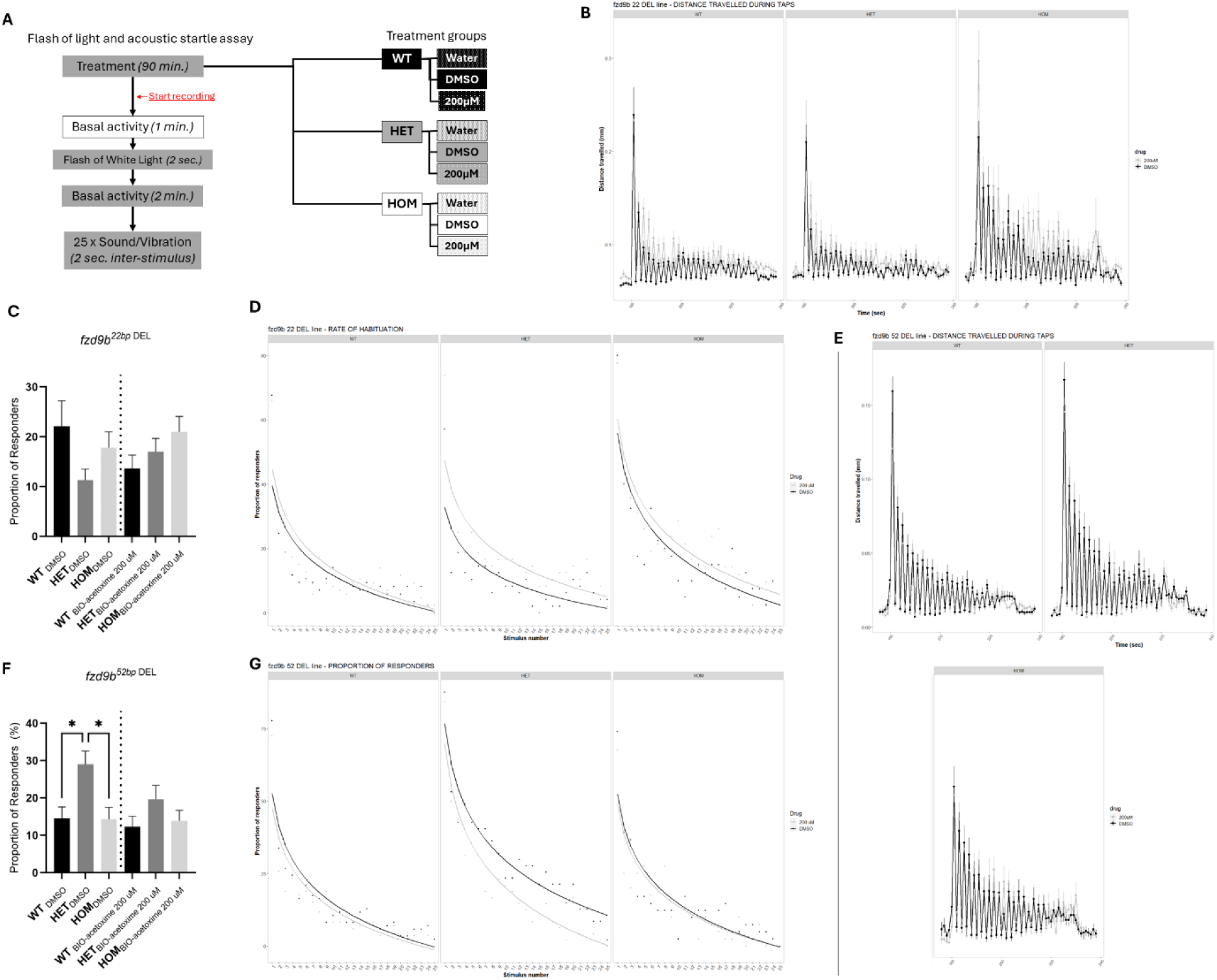
Flash of light and acoustic startle assay in 5 dpf zebrafish following exposure to BIO-acetoxime. **A)** Diagram of the assay and treatment groups. After 90-minutes exposure to BIO-acetoxime in the dark, zebrafish were recorded for 1 minute to assess basal activity also in darkness. Then, subjects were exposed to a 2-second illumination with white light and back to dark for 2 minutes. Finally, zebrafish were subjected to 25 consecutive sound/vibration stimuli with an inter-stimulus interval of 2 seconds also in darkness. Treatment conditions included: WT fish treated with water (WT-water, control), WT fish treated with DMSO (WT-DMSO, vehicle control), WT fish treated with 200 μM of BIO-acetoxime (WT-200 μM) and both mutants HETs and HOMs treated similarly (HET-Water, HET-DMSO, HET-200μM, HOM-Water, HOM-DMSO, and HOM-200μM). Panels **B)**, **C)** and **D)** show distance travelled during taps, absolute proportion of responders to the taps and rate of habituation to the taps respectively for the *fzd9b^22bpDEL^* line in the presence of the drug-vehicle. Panels **E)**, **F)** and **G)** show similar findings for the *fzd9b^52bpDEL^* line. In both cases: * p < 0.05 vs. corresponding contrast.

In the *fzd9b^22bpDEL^* line, similarly to previous findings, there were no differences across genotypes in the basal locomotion before the flash of light and before the acoustic stimuli (Supplementary Table 3). Likewise, there were no differences across genotypes in the magnitude of response to the flash of light, nor in the recovery slope (Supplementary Table 3). In contrast with previous findings, after DMSO exposure, during the tap events, we observed a significant main effect of genotype and a significant two-way interaction between genotype and time on distance travelled such that HOMs travelled greater distances than WT (p < 0.05, Supplementary Table 3). When assessing the absolute proportion of responders, there were no significant differences between genotypes (p > 0.05). However, as seen previously, when assessing the rate of habituation over time, there was a significant main effect of genotype and a significant two-way interaction between stimulus events and genotype (see Supplementary Table 3), whereby HOM fish habituate more slowly than WT (p < 0.0001) and HET (p < 0.0001), while there were no significant differences between WT and HET (p > 0.05).

Treatment with BIO-acetoxime (200 µM), had no significant effect on basal locomotion, and no significant effect on the response to the flash of light (Supplementary Table 3). When assessing the overall distance travelled during tap events, there were no significant differences between treated and control fish within the same genotype (p > 0.05). However, there was a significant three-way interaction between genotype * drug * time (Supplementary Table 3), such that HOM treated with BIO-acetoxime travelled greater distances than WT_BIO-acetoxime_ (p < 0.05) and HET_BIO-acetoxime_ (p < 0.05; Fig. 9B). As for the proportion of responders, BIO-acetoxime had no effect on absolute proportion of responders (p > 0.05; Fig. 9C). When analysing the rate of habituation over time, as seen previously, there was a main effect of genotype such that HOM habituate more slowly. There is a main effect of treatment such that fish treated with BIO-acetoxime habituate more slowly than controls (p < 0.0001), and there was a significant two-way interaction between genotype * drug, and a significant three-way interaction between genotype * drug * stimulus event (Supplementary Table 3), whereby HOM treated with BIO-acetoxime habituate more slowly than WT_BIO-acetoxime_ and HET_BIO-acetoxime_ (p < 0.0001 and p < 0.0001 respectively; Fig. 9D).

In the *fzd9b^52bpDEL^* line, as previously, there were no differences in baseline locomotion across genotypes, nor in the distance travelled during acoustic stimuli (Supplementary Table 3). Similarly, there were no differences across genotypes in the magnitude of response to the flash of light, nor in the recovery slope (Supplementary Table 3). During the tap events, there was no significant effect of genotype on distance travelled (Supplementary Table 3). When assessing the absolute proportion of responders, as seen before, there was a significant main effect of genotype on the absolute proportion of responders (Supplementary Table 3), such that a higher percentage of HET fish responded to the acoustic stimuli compared to WT (p < 0.05) and HOMs (p < 0.05) siblings, while there were no differences between WTs and HOMs (p > 0.05). Regarding the rate of habituation over time, there was a significant two-way interaction between stimulus events and genotype (Supplementary Table 3), whereby HET fish habituate more slowly than WT and (p < 0.0001) and HOM (p < 0.0001), and no differences between WT and HOM (p > 0.05).

As for the *fzd9b^22bp^ ^DEL^* line, exposure of *fzd9b^52bpDEL^* to BIO-acetoxime (200 µM), had no significant effect on locomotion before the flash of light and before the taps, no significant effect on the response to the flash of light, and no significant main effect in the distance travelled during the tap events (p > 0.05; Fig. 9E; Supplementary Table 3). Regarding the absolute proportion of responders, there was no main effect of BIO-acetoxime and no genotype by drug interaction (p > 0.05; Fig. 9F). When analysing the rate of habituation over time, in the presence of drug, in contrast to results seen in the *fzd9b^22bp^ ^DEL^* line, there was a main effect of BIO-acetoxime (Supplementary Table 3) whereby treated fish habituate more quickly than controls (p < 0.0001), and there was a significant three-way interaction between genotype * drug * stimulus number (Supplementary Table 3), whereby HET treated with BIO-acetoxime habituate more slowly than WT_BIO-acetoxime_ (p < 0.0001) and HOM_BIO-acetoxime_ (p < 0.0001; Fig. 9G).

## Discussion

Here, we have generated two mutant lines with disrupted *fzd9b* (*fzd9b^22bpDEL^* and *fzd9b^52bpDEL^* lines) to test the hypothesis that disruption of Fzd9b signalling contributes to altered stress reactivity and social behaviour in zebrafish as seen in WS patients. We generated loss-of-function zebrafish and confirmed disruption of Fzd9b signalling using expression analysis of genes known to be regulated by Fzd9. Behavioural analysis demonstrated that *fzd9b* mutant fish have altered stress- and anxiety-related responses at larval and adult stages. Additionally, we found that sociability, which is increased in WS individuals, was marginally increased in the heterozygous fish from the *fzd9b^52bpDEL^* line. Finally, we attempted to pharmacologically rescue their stressful phenotype via an acute manipulation of the Wnt/β-catenin pathway. However, in neither line were behaviour effects reversed by treatment with antagonists of the canonical Wnt signalling pathway. Our data suggests that *FZD9* contributes to the altered stress- and anxiety-phenotypes reported in WS but argues against Wnt canonical signalling as a potential pharmacological intervention to tackle this WS-associated recurrent neurological feature.

### Disrupting *fzd9b* leads to compensatory upregulation of *fzd9b*, *wnt5b* and *tafa5l*

In zebrafish, *fzd9b* is a single exon gene and, therefore, as expected, the mRNA surveillance pathway did not lead to a decay in its expression. Thus, both versions of the mutant mRNA are predicted to remain in heterozygotes, and, accordingly, both complete and truncated proteins are predicted to be generated. Conversely, only the truncated protein is predicted to remain in the homozygous mutants.

The predicted proteins generated by the mutant mRNAs differ in the two lines: in *fzd9b^22bpDEL^* the mutant mRNA generates a 24 aa polypeptide that consists of the first 20 aa of the mature Fzd9b protein plus 4 mis-matched aa. The *fzd9b^52bpDEL^* protein consists of the first 23 aa of the mature Fzd9b protein plus 44 mis-matched aa. Neither of these truncated proteins contains the intracellular domain required for their activation and downstream signalling (Agostino, Pohl and Dharmarajan, 2017) suggesting they act as loss-of-function mutations. However, interestingly, the *fzd9b^52bpDEL^* protein includes 3 cysteines in the mis-matched domain which might increase its binding affinity to the cysteine-rich homology domain found in the secreted frizzled related protein 2, Sfrp2, a member of the Sfrp family that binds to Wnt ligands and Frizzled receptors to modulate their function (Ladher *et al*., 2000; Agostino, Pohl and Dharmarajan, 2017). Although more studies are necessary, it is possible that the additional aas enable this *fzd9b^52bpDEL^*peptide to interact with Wnt proteins to disrupt signalling through other receptors.

Furthermore, the qPCR analysis revealed an increased production of *fzd9b* mRNA in the HETs of both mutant lines that was no longer present in the HOMs. This increased amount of *fzd9b* mRNA seen in HETs suggests that a reduction in functional Fzd9b led to a compensatory increase in *fzd9b* expression that was not seen on the complete abolition of functional *fzd9b* mRNA. A similar increase in the express of *wnt5b*, a ligand for Fzd9b, but not of *wnt5a*, was also seen in mutant from both lines, although it only reached significance in HETs from the *fzd9b^52bpDEL^*line and in HOMs from the *fzd9b^22bpDEL^* line. In rodents, WNT5A and FZD9 interact to mediate an increase in density of dendritic spines (Ramírez *et al*., 2016). Our data suggest that in zebrafish, contrary to rodents, Wnt5b instead of Wnt5a may be a key interacting partner for Fzd9b in this regard. Interestingly, the number and size of dendritic spines is significantly increased in WS individuals, which may be related to an adaptive up-regulation of Fzd9/Wnt signalling and contribute to functional alterations associated with this syndrome (Chailangkarn *et al*., 2016).

We also saw an apparent increase in *wnt9a* expression, the Fzd9b cognate ligand involved in hematopoietic expansion in zebrafish (Grainger *et al*., 2019), in the *fzd9b^22bpDEL^* line, but this did not reach significance. Nonetheless, the up regulation of expression of components of the Fzd9 signalling pathway seen here (*fzd9b*, *wnt5a*) is consistent with a reduction in functional protein as predicted by the mutations carried by the generated lines. We may speculate that, on reduction of Fzd9b signalling alternative compensatory mechanisms are induced, perhaps involving other Fzd receptors such as Fzd9a, the other paralogue for this gene in zebrafish (Nikaido, Law and Kelsh, 2013; Diss *et al*., 2014). The increase in expression of *fzd9b* and *wnt5b* suggests a compensatory increase in downstream components of Fzd9b/Wnt signalling.

To further test that the mutated mRNA led to reduced Fzd9b function we examined the expression level of *cmyb* by *in situ* hybridization*. cmyb* is a marker of HPSC expansion the expression of which is positively regulated by Fzd9b (Grainger *et al*., 2019). As expected, we saw a decrease in *cymb* expression domain in HETs which was further decreased in HOMs, for both lines. This agrees with previous findings using either a morpholino approach or targeted CRISPR/cas9 editing in hematopoietic endothelium to knockdown fzd9b expression (Grainger *et al*., 2019).

We also evaluated the expression of zebrafish homologues of 2 genes, *tafa5l* (also named *fam19A5*) and *scn4aa7b,* that are differentially expressed in neurons derived from WS patients compared to control individuals, but unaffected in derived neurons from WS patients carrying deletions that spare the *FZD9* gene (Chailangkarn *et al*., 2016). As in derived neurons from WS patients that are heterozygous for the *FZD9* gene deletion, we saw an increase in expression of *tafa5l* in the HOMs from the *fzd9b^22bpDEL^* line and in both HETs and HOMs from the *fzd9b^52bpDEL^* line. Thus, similar to humans, the expression of *tafa5l* seems to be regulated by Fzd9b in zebrafish. *tafa5l* codes for a chemokine-like protein involved in cell proliferation and migration via G protein-coupled receptors. Expression of this protein seems to be restricted to brain structures (Tom Tang *et al*., 2004) and has been associated with, among others, cognitive impairment in vascular dementia (Li *et al*., 2020).

Additionally, in *fzd9b^52bpDEL^* heterozygotes, an increase in the expression of *scn4aa* and *scn4ab* was also seen. This contrasts with findings in neurons derived from iPSC from WS patients where there is a significant decrease in *SCN4a* expression (Chailangkarn *et al*., 2016). However, the study conducted by Chailangkam examined expression in isolated neurons derived from iPSCs, whereas our study examined whole body expression. As *scn4a* is strongly expressed in muscle, the differences in findings may be related to the tissue sample. In zebrafish, there are two paralogs, *scn4aa* and *scn4ab*, which code for two voltage-gated Na^+^ channel α-subunits. From the two, *scn4ab* seems the one that retained the original function of propagating transmembrane currents in skeletal muscle in many teleost (Arnegard *et al*., 2010). Nonetheless, an increase in muscle expression may mask any decrease in neuronal expression of *scn4a* homologues in zebrafish. Importantly, musculoskeletal abnormalities are commonly seen in WS individuals (Copes, Pober and Terilli, 2016) which may be related to altered *SCN4a* expression.

### Disrupting *fzd9b* in zebrafish results in a marginal effect on sociability and altered stress- and anxiety-like responses at larval and adult stages respectively

Pro-social behaviour is a key feature of WS patients. Thus, we tested whether *fzd9b* participated in the early social behaviour of zebrafish. Wnt signalling has been shown to regulate neurodevelopment through its non-canonical pathway (Mayor and Theveneau, 2014) and, additionally, inhibition of GSKb (as occurs on activation of the canonical pathway) is associated with pro-social behaviour in rodents (Latapy *et al*., 2012). Interestingly, only HETs from the *fzd9b^52bpDEL^* line showed a significantly higher predisposition to interact with multiple conspecifics than WT fish. Although further studies are required to understand this finding, the increase in sociability may be related to the up regulation in the expression of *fzd9b* seen in HETs or, perhaps, to the mutant protein interacting with other Wnt pathway proteins.

However, as no effect on social behaviour was seen in mutants from the other *fzd9b* line, where there was also an up-regulation in *fzd9b* mRNA, loss of *fzd9b per se* may make a minor contribution to the increased sociability seen in WS. Nonetheless, this mild change in sociability, together with the stress- and anxiety-like alterations reported here, might be supporting further the interaction between sociability and anxiety reported in WS (Ng-Cordell *et al*., 2018); although, as our data suggests, *FZD9* might not be a top contributor to this interaction. We and others have recently pointed that *BAZ1B*, one of the other genes deleted in classical WS, could be playing a key role in the ontogeny of social behaviour through regulation of maturation of the neural crest (Lalli *et al*., 2016; Zanella *et al*., 2019; Torres-Pérez *et al*., 2023). Nonetheless, and despite variation between lines, our behavioural data strongly points at *fzd9b* as a major contributor to altered stress- and anxiety-like responses in larval and adult zebrafish respectively.

Stress reactivity in the larvae was assessed using the FLDT paradigm and the acoustic startle response. FLDT assay is an established test to assess differences in stress-reactivity in zebrafish (Peng *et al*., 2016; Lee *et al*., 2019; García-González *et al*., 2021; Faught and Vijayan, 2022). In WT 5 dpf larvae, on change of light conditions from dark to light, the fish exhibit a freezing response with the extent of the decrease in locomotion and the rate of recovery being indicative of degree of stress reactivity/ anxiety. Similarly, on change from light to dark, the fish exhibit a rapid increase in locomotion followed by gradual reduction to baseline. Again, the magnitude of the increase in locomotion and the rate of recovery are taken as indicative of stress reactivity (García-González *et al*., 2021). In both lines, mutant larvae demonstrated similar recovery to the changes in illumination compared to wildtype siblings indicating no differences in stress-reactivity due to forced changes in environmental light.

Nonetheless, upon exposure to repeated acoustic startle, we found differences suggesting increased stress-reactivity in the mutants. The repeated acoustic startle assay is primarily used to study startle and habituation responses to sudden sound stimuli, which can be associated with alterations in the fish’s stress response (Lee *et al*., 2019; Torres-Pérez *et al*., 2023). Accordingly, HOMs from the *fzd9b^22bpDEL^* and HETs from the *fzd9b^52bpDEL^* lines took longer to habituate. Although the startle response is a complex behaviour influenced by various factors, a longer time to habituate (or lack of habituation) indicates increased sensitivity to stressors. Interestingly, hypersensitivity to sound (hyperacusis) is a common feature in WS (Pérez Jurado *et al*., 1996), and increased auditory startle reflex, associated with autonomic hyper-reactivity, is a prevalent characteristic in children with anxiety disorders (Bakker *et al*., 2009). Therefore, this longer rate of habituation at zebrafish’s larval stages, in combination with the output of our behavioural assay in the adults, suggests that mutants have altered anxiety-like responses.

Anxiety-like behaviours during adulthood were assessed by the tank diving assay. In this assay, the anxiety-like response is measured as the time and distance fish spent near the bottom of the tank (Evans *et al*., 2021). Although the phenotype was surprisingly different between the lines, mutants from both lines showed altered anxiety-like behaviour in adulthood: both HETs and HOMs from the *fzd9b^22bpDEL^* line spent overall more time close to the bottom, which is a sign of an increased anxious-like phenotype. In contrast, HETs from the *fzd9b^52bpDEL^* line spent less time in the bottom third, an established sign of reduced anxiety-like behaviour (Egan *et al*., 2009). HETs from the *fzd9b^52bpDEL^* line also had more visits to the top third of the tank, which again is attributed to a lower anxiety-like response. We have no solid explanation for this finding. It may relate to fish being erratic, rather than anxious-like but neither freezing nor darting rates were different between genotypes (data not shown). Nonetheless, the increased total distance move of this HETs might be suggesting such phenotype. Similar discrepancies in anxiety-like behaviours between lines, as measured in the novel tank diving assay, have been found in zebrafish following early-life dysregulation of the HPI axis (Eachus, Choi and Ryu, 2021). Differences between the *fzd9b^22bpDEL^* and *fzd9b^52bpDEL^* lines may, again, relate to the differences in the truncated proteins generated.

### Altered stress reactivity is not mediated by Wnt canonical pathway in the *fz9b* mutant larvae

As our behavioural analysis gave some support for altered anxiety-like responses in *fzd9b* loss-of-function lines and Wnt signalling has been suggested to modulate anxiety-related behaviour through the canonical signalling pathway (Latapy *et al*., 2012; Korem *et al*., 2017), we examined the ability of an acute treatment aiming at activating this pathway to reverse effects on stress related behaviour in the two lines. We used BIO-acetoxime, a drug that selectively inhibits GSK-3 (Aourz *et al*., 2019), which has a central role in the canonical Wnt/β-catenin signalling pathway.

An acute treatment with BIO-acetoxime was not able to restore the behaviour of the mutant fish to the levels of WTs in either of the two lines studied. This behaviour was evaluated with an assay combining a flash of light and acoustic startle, a test known to rely on HPI reactivity (Lee *et al*., 2019; Torres-Pérez *et al*., 2023), following 90 minutes of exposure to the drug. Treatment with 200 µM BIO-acetoxime was not able to restore the habituation to the acoustic startle.

The observed differences in stress reactivity and habituation in the non-treated animals were similar to those reported in the previous larval assays, thus differences were maintained in the mutants regardless of treatment. Therefore, altered stress reactivity prevailed in the mutants despite the treatment.

Taking all together, our data suggest that lack of functional *fzd9b* influences stress-reactivity in larval zebrafish. However, targeting the canonical pathway from the Wnt/β-catenin signalling system does not have an impact on the observed differences in stress reactivity in the fish with disrupted *fzd9b*.

## Conclusion and final remarks

Variations in the dosage of functional *fzd9b* in zebrafish resulted in changes at both molecular and behavioural levels. In the mutant fish, the stress- and anxiety-driven responses, which are affected in WS individuals with a copy of this gene missing, were significantly altered. Additionally, acute treatment aiming to restore the Wnt/β-catenin pathway was not able to revert those behavioural manifestations in larval zebrafish, thus suggesting that this endophenotype is regulated by a non-canonical signalling cascade. Nonetheless, our data on *fzd9b*’s influence of both stress and anxiety responses suggests that future studies aiming to target the non-canonical Wnt signalling pathway, or FZD9 itself, could be of therapeutic use to tackle altered stress-reactivity in WS and associated disorders. In this regard, long-term treatment with Foxy 5, a Wnt5a-derived peptide currently in clinical trials for metastatic cancers (Xu *et al*., 2020) and known to influence synaptic plasticity in mice (Vargas, Fuenzalida and Inestrosa, 2014), could be of interest.

This research also shows that, although stress and anxiety responses are altered, lack of functional *fzd9b* does not have a major effect on sociability, another feature associated with WS. Recently, we have demonstrated that *baz1b*, another gene missing in classical WS, regulates the ontogeny of sociability in zebrafish (Torres-Pérez *et al*., 2023). Additionally, our previous research suggests that lack of *baz1b* leads to reduced stress- and anxiety-related behaviours via developmental alterations of the neural crest in zebrafish. Both *baz1b* and *fzd9b*, each with an opposing effect on stress- and anxiety-related phenotypes, are missing in classical WS, which could account for the varying prevalence of stress- and anxiety-related behaviours in WS. Differential penetrance for these two genes, alone or in combination with other genetic and environmental factors (e.g. early life stress), could be determining the final state of the HPA/HPI axis. Thus, our research highlights the need to establish specific gene-phenotype correlations, of particular importance in disorders spanning multiple genes such as WS.

Our study is not without limitations: At a molecular level, we observed an upregulation of *fzd9b* in the heterozygous fish. We also provided evidence that, contrary to rodents, Wnt5b, instead of Wnt5a, interacts with Fzd9b. Further, in agreement with human data, the expression of *tafa5l* was affected by the functional levels of Fzd9b. However, the molecular mechanism(s) leading to transcriptional upregulation of *fzd9b* itself and the associated genes, as well as the specific function(s) and targeted tissue/organs, were not assessed. Additionally, although our study proposes that *fzd9b* contributes to the stress- and anxiety-alterations associated while having little or no impact on their social abilities, an interplay between anxiety and social functioning (driven by *FZD9*-related genetic networks or independently) might still persist in WS (Riby *et al*., 2014; Willfors *et al*., 2021). Future investigations attempting to assess the combined anxiety and social phenotypes might elucidate the existence of synergies.

## Materials and Methods

### Animal husbandry

All procedures involving live animals were reviewed and approved by the Queen Mary University of London ethics committee (AWERB) following consultation of the ARRIVE guidelines (NC3Rs, UK) and conducted in accordance with the Animals (Scientific Procedures) Act, 1986 and Home Office Licenses.

Zebrafish used in this study originated from a Tübingen background line. Subjects were kept at ∼25–28 °C in a system of recirculating water and at a light:dark cycle of 14:10. Fish were fed two times a day with dry food (ZM-400, Zebrafish Management Ltd) or live brine shrimp (*Artemia salina*). Breeding and rearing of the offspring were done similarly as reported previously (Torres-Pérez et al., 2023).

### Generation of *fzd9b* loss-of-function (LoF) zebrafish

We used a similar CRISPR/Cas9 strategy to induce the LoF to that previously reported in our group (Torres-Pérez *et al*., 2023) but tailored to the zebrafish gene *fzd9b* (Ensembl gene ID: ENSDARG00000014673). The crRNA (5’ACAGCCAUACUCCACCAGAG) was designed to overlap a restriction site for Bsl1 including a PAM site and located within the upstream (5’) half portion of this single exon gene. Primers used for genotyping: forward 5’AGGGCATCGGATACAACCTC and reverse 5’TTCAAAGATTCAGGCCAGGC (Fig. 1A). Zebrafish lines can be requested to the corresponding authors.

### Quantitative PCR (qPCR)

At 5 days post-fertilization (dpf), zebrafish larvae from visually similar developmental stages were collected in groups of 16 individuals from all corresponding genotypes (N = 4-5 per genotype) and stored in RNAlater (Thermo Fisher) until use. RNA extraction, cDNA synthesis and quantitative PCR (qPCR) were performed and validated as previously reported (Torres-Pérez et al., 2023). Two housekeeping genes (*ef1a* and *rpll3α*) were used to normalise the data and assess relative expression changes for the different genes studied. Gene names, sequence of the primers and additional details can be found in Supplementary Table 1. Primers for *fzd9b* were designed to align an upstream location from the mutation site used to generate the LoF.

### Whole body *in situ* hybridization (ISH) and % of area stained

After carefully staging the zebrafish embryos and selecting for no visual differences in development, embryos from all genotypes (N = 7 per genotype) were collected at 40 hpf and fixed with 4% paraformaldehyde (PFA; Sigma, Gillingham, UK) in phosphate buffered saline (PBS) overnight. Following day, fixed embryos were rinsed in PBS supplemented with Tween (0.05% v/v), dehydrated in ascending methanol series (25%, 50%, 70%, 80%, 90%, 100% methanol, 5 min each), and stored in 100% methanol and kept at -20 °C until use. The procedure for *in situ* hybridization (ISH) was similar to that used previously (Torres-Pérez *et al*., 2023). Briefly, RNA probes against *cmyb* (ENSDART00000075726.5) were generated and amplified by PCR from cDNA generated using following set of primers: Forward 5’CTGCTAAAGTCAGCCCAACTCC and Reverse 5’ ATCGTCTGCTCTTCCGTCTTCC. Lateral view pictures of the stained zebrafish were taken using a Leica MZ75 microscope.

Fiji (Schindelin *et al*., 2012) was used to measure the percentage of positive staining at their tails. First, if required, pictures were rotated until tails were in the horizontal plane. Then, pictures were converted to greyscale (8-bit), histogram equalised and resulting images transformed to binary using the Phasalkar’s method (Phansalkar *et al*., 2011). A rectangular region of interest (ROI) was drawn with a width extending from the end of the ‘yolk sack’ to the end of the ‘yolk sack extension’ and with the minimal height required to include the embryo’s tail within that segment. The percentage of positive area from the ROI was measured and used for analysis.

## Behavioural experimental design

### Experiment 1: Sociability in 3 weeks old zebrafish

To assess <u>social behaviours in young zebrafish</u>, we performed this assay as originally developed by Dreosti et al. 2015 (Dreosti *et al*., 2015) with similar modifications as described in our previous study (Torres-Pérez *et al*., 2023), but only at three weeks of age (between 20 to 22 dpf). Briefly, fish were tested between 10 am and 7 pm, with each test subject matched to stimuli fish from the same genotype and with similar age/size. Experiments were performed simultaneously in two DanioVision Observation Chambers with a minimum of two independent experiments conducted per day and balanced sides of stimuli presentation. Total motility and position within the arena were recorded and analysed using EthoVision XT software and data outputted in 15-minute time bins. Both Social Preference Index (SPI) and Correlation Index (r = SPI_ExperimentalPhase_ – SPI_AcclimationPeriod_) were calculated as described previously (Torres-Pérez *et al*., 2023).

### Experiment 2: Forced changes in environmental light at 5 dpf

#### Forced light-dark transition (FLDT) at 5 dpf

Test performed with similar conditions as previously described (Glazer *et al*., 2018; Torres-Pérez *et al*., 2023) and used to to assess stress-related behaviour in zebrafish at 5 days post-fertilization (dpf). Briefly, singly housed 5 dpf larvae were recorded in the dark (infrared conditions) for 10 minutes as baseline. This period was followed by 3 cycles of light/dark transition cycles, each of 10 minutes light followed by 10 minutes darkness. Distance travelled was recorded with the Ethovision XT software (Noldus Information Technology, Wageningen, NL) and data exported in 1 min and 1 sec time bins. Tests were conducted in parallel, between 9 am and 4 pm and during three consecutive days with two independent experiments per day. To ensure experimenter was blind to genotype, these assays were done using the offspring of in-crossing multiple zebrafish heterozygous for the corresponding mutation (*fzd9b^+/-^*x *fzd9b^+/-^*). Genotyping (as explained above) was carried out following each behavioural procedure.

### Experiment 3: Acoustic startle habituation at 5 dpf

#### Flash of light and repeated acoustic startle

This test was performed in 5 dpf larvae placed in individual wells (24-multiwell plates) darkness. After an initial 10 min period of dark (baseline), larvae were exposed to a 2-second flash of light, followed by 1 min recovery. Subsequently, larvae were exposed 20 sound/vibration consecutive stimuli with an inter-stimulus period of 2 seconds and an additional 2-second flash of light. Tests were conducted between 9 a.m. and 4 p.m. Distance travelled was similarly recorded and data outputted as with the FLDT assay (Experiment 2).

### Experiment 4: Anxiety-like phenotypes in adult zebrafish

#### Novel tank diving assay

This assay is used in zebrafish to assess anxiety-driven responses. It was performed similarly as previously described in adult zebrafish (Evans *et al*., 2021). To avoid ‘tank effect’, at least two tanks of zebrafish from each genotype (WT, HET and HOM) were used. 3-month-old zebrafish from the different genotypes were tested simultaneously between 9 am and 2 pm. Zebrafish were individually placed in a trapezoid tank (novel shape tank) containing fresh fish system water (water replaced every 2-3 trials) and side-view recorded for 6 minutes. Distance to the bottom and total motility were tracked and analysed with EthoVision software. Data was outputted in time bins of 30 seconds.

### Experiment 5: Inhibiting GSK3 in the flash of light and repeated acoustic startle assay

For this experiment (Fig. 9A), the genotype of the 5 dpf zebrafish tested was known as they needed to be allocated to corresponding conditions within each 48-well plate. Each plate had at least 6 different conditions and around 4-6 replicates for each condition. The position of the different conditions/replicates was randomly allocated between plates. Zebrafish were the result of either HOM or WT in-crosses or HOM x WT crossings. Multiple zebrafish from all breeding combinations were crossed simultaneously and the offspring reared in multiple plates and similar incubation conditions until test.

5 dpf zebrafish from both mutant genotypes, HET and HOM, were singly housed and subjected to 90-minute incubations with 6-bromoindirubin-3’-oxime (BIO-acetoxime; HY-15356, Cambridge Bioscience) inside a dark DanioVision Chamber. A dose of 200 μM of BIO-acetoxime was used as previous toxicological evaluation done by Aourz et al. 2019 demonstrated this to be maximal tolerated concentration for larval zebrafish without inducing any locomotor alterations (Aourz *et al*., 2019). Intra-genotype controls included individuals treated with either water or the drug vehicle dimethyl sulfoxide (DMSO). Additional controls were WT zebrafish treated with either water, DMSO or the highest dose of BIO-acetoxime (200 μM). Therefore, experiment consisted in 9 conditions (Fig. 9) as follow: 1) WT water, 2) WT DMSO, 3) WT 200 μM, 4) HET Water, 5) HET DMSO, 6) HET 200 μM, 7) HOM water, 8) HOM DMSO, 9) HOM 200 μM.

#### Flash of light and acoustic startle (Fig. 9)

After 90 minutes exposure to the drug, zebrafish were subjected to a behavioural test that combined a light-locomotor behavioural assay and a startle tap test habituation assay similar to previously described (Torres-Pérez *et al*., 2023). Briefly, protocol consisted in 1 minutes of basal activity in darkness, 2-second illumination with white light, back to dark for 2 minutes and then 25 consecutive sound/vibration stimuli with an inter-stimulus interval of 2 seconds. This combined assay was used instead of the previously mentioned test (FLDT or Acoustic startle habituation) to be able to accommodate all conditions within similar testing hours (from 9 am to 2 pm). Ethovision XT software was used to record distance travelled and data were outputted in one-second time bins.

### Statistics

Molecular data were analysed with GraphPad Prism 9.0.2 for Windows (GraphPad Software, San Diego, California USA, www.graphpad.com). Each dataset was visually assessed for normal distribution with the homoscedasticity and the normal QQ plots. If normality assumptions were violated, a non-parametric test was used. Data from qPCR analysis was analysed using a two-way repeated measures ANOVA with “genotype” and “time” as factors and “subject” as matched set and using Bonferroni’s multiple comparison test. Data from the % of tail *cmyb*+ was analysed using the non-parametric Kruskal-Wallis test (unpaired samples) using Dunn’s test to correct for multiple comparisons. All effects are reported significant at p < 0.05.

Data from all larval behavioural assays (5 dpf) and novel tank assay were analysed similarly as our previous publication (Torres-Pérez *et al*., 2023). Briefly, we used RStudio (version 1.2.5042) and results reported with respect to a type-1 error rate of α = 0.05. R packages used include: ‘lme4’ with, if applicable, a post-hoc Tukey test using the R package ‘multcomp’. Percentage of fish responding to stimulus was analysed with GraphPad Prism 9.0.2 using Anova with repeat measures to look at interaction between total number of responders in all TAP events and genotype as explanatory variables. When applicable, Tukey test was used to correct for multiple comparisons.

Data from the sociability assay was analysed using GraphPad Prism 9.0.2 for Windows similarly as previous described (Torres-Pérez *et al*., 2023). In short, a two-tailed Wilcoxon test to assess changes in SPI in each condition (genotype, contrast) and a Kruskal-Wallis test with repeated measures and applying Dunn’s multiple comparisons test for correlation index. Statistical significance was defined at p < 0.05.

## Acknowledgments

We are really thankful to Mr Luca Galantini for his support, assistance and guidance at QMUL’s zebrafish facility.

## Data availability

Generated data can be found at https://zenodo.org/ (doi: 10.5281/zenodo.10548489). Data will be publicly available after acceptance for publication.

## Author contributions statement

**JVTP**: designed the experiments, performed research, data analysis and wrote the manuscript. **AL:** performed research, data analysis and contributed to writing of the manuscript. **AMM**: data analysis and contributed to writing of the manuscript. **SA**: performed research. **WH:** analysed data. **CHB**: supervised research, obtained funding for this research and writing the manuscript. All authors read and approved the final manuscript.

## References

de Abreu, M.S. et al. (2021) ‘Understanding how stress responses and stress-related behaviors have evolved in zebrafish and mammals’, Neurobiology of Stress, 15. Available at: 10.1016/j.ynstr.2021.100405.

Agostino, M., Pohl, S.Ö.G. and Dharmarajan, A. (2017) ‘Structure-based prediction of Wnt binding affinities for frizzled-type cysteine-rich domains’, Journal of Biological Chemistry, 292(27). Available at: 10.1074/jbc.M117.786269.

Aourz, N. et al. (2019) ‘Identification of GSK-3 as a Potential Therapeutic Entry Point for Epilepsy’, ACS Chemical Neuroscience, 10(4), pp. 1992–2003. Available at: 10.1021/acschemneuro.8b00281.

Arnegard, M.E. et al. (2010) ‘Old gene duplication facilitates origin and diversification of an innovative communication system - Twice’, Proceedings of the National Academy of Sciences of the United States of America, 107(51). Available at: 10.1073/pnas.1011803107.

Bakker, M.J. et al. (2009) ‘Increased whole-body auditory startle reflex and autonomic reactivity in children with anxiety disorders’, Journal of Psychiatry and Neuroscience, 34(4).

Basnet, R.M. et al. (2019) ‘Zebrafish larvae as a behavioral model in neuropharmacology’, Biomedicines. Available at: 10.3390/BIOMEDICINES7010023.

Beppi, C., Straumann, D. and Bögli, S.Y. (2021) ‘A model-based quantification of startle reflex habituation in larval zebrafish’, Scientific Reports, 11(1). Available at: 10.1038/s41598-020-79923-6.

Beurel, E., Grieco, S.F. and Jope, R.S. (2015) ‘Glycogen synthase kinase-3 (GSK3): Regulation, actions, and diseases’, Pharmacology and Therapeutics. Available at: 10.1016/j.pharmthera.2014.11.016.

Bradford, Y.M. et al. (2017) ‘Zebrafish models of human disease: Gaining insight into human disease at ZFIN’, ILAR Journal, 58(1), pp. 4–16. Available at: 10.1093/ilar/ilw040.

Chailangkarn, T. et al. (2016) ‘A human neurodevelopmental model for Williams syndrome’, Nature, 536(7616), pp. 338–343. Available at: 10.1038/nature19067.

Cohen, H. et al. (2006) ‘Blunted HPA Axis Response to Stress Influences Susceptibility to Posttraumatic Stress Response in Rats’, Biological Psychiatry, 59(12). Available at: 10.1016/j.biopsych.2005.12.003.

Copes, L.E., Pober, B.R. and Terilli, C.A. (2016) ‘Description of common musculoskeletal findings in Williams Syndrome and implications for therapies’, Clinical Anatomy. Available at: 10.1002/ca.22685.

Diss, G. et al. (2014) ‘Molecular mechanisms of paralogous compensation and the robustness of cellular networks’, Journal of Experimental Zoology Part B: Molecular and Developmental Evolution. Available at: 10.1002/jez.b.22555.

Dreosti, E. et al. (2015) ‘Development of social behavior in young zebrafish’, Frontiers in Neural Circuits, 9(AUGUST). Available at: 10.3389/fncir.2015.00039.

Eachus, H., Choi, M.K. and Ryu, S. (2021) ‘The Effects of Early Life Stress on the Brain and Behaviour: Insights From Zebrafish Models’, Frontiers in Cell and Developmental Biology. Available at: 10.3389/fcell.2021.657591.

Egan, R.J. et al. (2009) ‘Understanding behavioral and physiological phenotypes of stress and anxiety in zebrafish’, Behavioural Brain Research, 205(1). Available at: 10.1016/j.bbr.2009.06.022.

Evans, J.R. et al. (2021) ‘Stress reactivity elicits a tissue-specific reduction in telomere length in aging zebrafish (Danio rerio)’, Scientific Reports, 11(1). Available at: 10.1038/s41598-020-79615-1.

Ewart, A.K. et al. (1993) ‘Hemizygosity at the elastin locus in a developmental disorder, Williams syndrome’, Nature Genetics, 5(1). Available at: 10.1038/ng0993-11.

Faught, E. and Vijayan, M.M. (2022) ‘Coordinated Action of Corticotropin-Releasing Hormone and Cortisol Shapes the Acute Stress-Induced Behavioural Response in Zebrafish’, Neuroendocrinology, 112(1). Available at: 10.1159/000514778.

Fusco, C. et al. (2014) ‘Smaller and larger deletions of the Williams Beuren syndrome region implicate genes involved in mild facial phenotype, epilepsy and autistic traits’, European Journal of Human Genetics, 22(1). Available at: 10.1038/ejhg.2013.101.

García-González, J., et al. (2021) ‘Behavioral effects of developmental exposure to jwh-018 in wild-type and disrupted in schizophrenia 1 (Disc1) mutant zebrafish’, Biomolecules, 11(2), pp. 1–23. Available at: 10.3390/biom11020319.

Glazer, L. et al. (2018) ‘Developmental Exposure to Low Concentrations of Organophosphate Flame Retardants Causes Life-Long Behavioral Alterations in Zebrafish’, Toxicological Sciences, 165(2). Available at: 10.1093/toxsci/kfy173.

Grainger, S. et al. (2019) ‘EGFR is required for Wnt9a–Fzd9b signalling specificity in haematopoietic stem cells’, Nature Cell Biology, 21(6), pp. 721–730. Available at: 10.1038/s41556-019-0330-5.

Järvinen, A., Ng, R. and Bellugi, U. (2015) ‘Autonomic response to approachability characteristics, approach behavior, and social functioning in Williams syndrome’, Neuropsychologia, 78. Available at: 10.1016/j.neuropsychologia.2015.10.012.

Korem, N. et al. (2017) ‘Role of beta-catenin and endocannabinoids in the nucleus accumbens in extinction in rats exposed to shock and reminders’, Neuroscience, 357. Available at: 10.1016/j.neuroscience.2017.06.015.

Ladher, R.K. et al. (2000) ‘Cloning and expression of the Wnt antagonists Sfrp-2 and Frzb during chick development’, Developmental Biology, 218(2). Available at: 10.1006/dbio.1999.9586.

Lalli, M.A. et al. (2016) ‘Haploinsufficiency of BAZ1B contributes to Williams syndrome through transcriptional dysregulation of neurodevelopmental pathways’, Human Molecular Genetics, 25(7). Available at: 10.1093/hmg/ddw010.

Latapy, C. et al. (2012) ‘Selective deletion of forebrain glycogen synthase kinase 3β reveals a central role in serotonin-sensitive anxiety and social behaviour’, Philosophical Transactions of the Royal Society B: Biological Sciences, 367(1601). Available at: 10.1098/rstb.2012.0094.

Lee, H.B. et al. (2019) ‘Novel zebrafish behavioral assay to identify modifiers of the rapid, nongenomic stress response’, *Genes*, Brain and Behavior, 18(2). Available at: 10.1111/gbb.12549.

Leggieri, A. et al. (2022) ‘Ankk1 Loss of Function Disrupts Dopaminergic Pathways in Zebrafish’, Frontiers in Neuroscience, 16. Available at: 10.3389/fnins.2022.794653.

Lense, M.D. and Dykens, E.M. (2013) ‘Cortisol reactivity and performance abilities in social situations in adults with williams syndrome’, American Journal on Intellectual and Developmental Disabilities, 118(5). Available at: 10.1352/1944-7558-118.5.381.

Li, J. et al. (2020) ‘Association of Serum FAM19A5 with Cognitive Impairment in Vascular Dementia’, Disease Markers, 2020. Available at: 10.1155/2020/8895900.

Martens, M.A., Wilson, S.J. and Reutens, D.C. (2008) ‘Research Review: Williams syndrome: A critical review of the cognitive, behavioral, and neuroanatomical phenotype’, Journal of Child Psychology and Psychiatry and Allied Disciplines. Available at: 10.1111/j.1469-7610.2008.01887.x.

Mayor, R. and Theveneau, E. (2014) ‘The role of the non-canonical Wnt-planar cell polarity pathway in neural crest migration’, Biochemical Journal. Available at: 10.1042/BJ20131182.

Morris, C.A. (1999) Williams Syndrome Synonym: Williams-Beuren Syndrome, Gene Reviews.

Ng-Cordell, E. et al. (2018) ‘Anxiety in Williams Syndrome: The Role of Social Behaviour, Executive Functions and Change Over Time’, Journal of Autism and Developmental Disorders, 48(3). Available at: 10.1007/s10803-017-3357-0.

Nicholson, P. and Mühlemann, O. (2010) ‘Cutting the nonsense: The degradation of PTC-containing mRNAs’, in Biochemical Society Transactions, pp. 1615–1620. Available at: 10.1042/BST0381615.

Nikaido, M., Law, E.W.P. and Kelsh, R.N. (2013) ‘A Systematic Survey of Expression and Function of Zebrafish frizzled Genes’, PLoS ONE, 8(1). Available at: 10.1371/journal.pone.0054833.

Noelanders, R. and Vleminckx, K. (2017) ‘How Wnt Signaling Builds the Brain: Bridging Development and Disease’, Neuroscientist. Available at: 10.1177/1073858416667270.

Okerlund, N.D. and Cheyette, B.N.R. (2011) ‘Synaptic Wnt signaling-a contributor to major psychiatric disorders?’, Journal of Neurodevelopmental Disorders. Available at: 10.1007/s11689-011-9083-6.

Peng, X. et al. (2016) ‘Anxiety-related behavioral responses of pentylenetetrazole-treated zebrafish larvae to light-dark transitions’, Pharmacology Biochemistry and Behavior, 145. Available at: 10.1016/j.pbb.2016.03.010.

Pérez Jurado, L.A., et al. (1996) ‘Molecular definition of the chromosome 7 deletion in Williams syndrome and parent-of-origin effects on growth’, American Journal of Human Genetics, 59(4).

Phansalkar, N. et al. (2011) ‘Adaptive local thresholding for detection of nuclei in diversity stained cytology images’, in *ICCSP 2011 - 2011 International Conference on Communications and Signal Processing*. Available at: 10.1109/ICCSP.2011.5739305.

Van Raay, T.J. et al. (2001) ‘Frizzled 9 is expressed in neural precursor cells in the developing neural tube’, Development Genes and Evolution, 211(8–9). Available at: 10.1007/s004270100174.

Ramírez, V.T. et al. (2016) ‘Wnt-5a/frizzled9 receptor signaling through the Gαo-Gβγ complex regulates dendritic spine formation’, Journal of Biological Chemistry, 291(36), pp. 19092–19107. Available at: 10.1074/jbc.M116.722132.

Riby, D.M. et al. (2014) ‘The interplay between anxiety and social functioning in Williams syndrome’, Journal of Autism and Developmental Disorders, 44(5). Available at: 10.1007/s10803-013-1984-7.

Royston, R. et al. (2021) ‘Anxiety characteristics in individuals with Williams syndrome’, Journal of Applied Research in Intellectual Disabilities, 34(4). Available at: 10.1111/jar.12864.

Schindelin, J. et al. (2012) ‘Fiji: An open-source platform for biological-image analysis’, Nature Methods. Available at: 10.1038/nmeth.2019.

Shah, S.M. et al. (2009) ‘Expression of Wnt receptors in adult spiral ganglion neurons: frizzled 9 localization at growth cones of regenerating neurites’, Neuroscience, 164(2). Available at: 10.1016/j.neuroscience.2009.08.049.

Stinton, C., Tomlinson, K. and Estes, Z. (2012) ‘Examining reports of mental health in adults with Williams syndrome’, Research in Developmental Disabilities, 33(1). Available at: 10.1016/j.ridd.2011.09.002.

Tanaka, M. et al. (2019) ‘Region- and time-dependent gene regulation in the amygdala and anterior cingulate cortex of a PTSD-like mouse model’, Molecular Brain, 12(1). Available at: 10.1186/s13041-019-0449-0.

Tom Tang, Y., et al. (2004) ‘TAFA: A novel secreted family with conserved cysteine residues and restricted expression in the brain’, Genomics, 83(4). Available at: 10.1016/j.ygeno.2003.10.006.

Torres-Pérez, J. V. et al. (2023) ‘baz1b loss-of-function in zebrafish produces phenotypic alterations consistent with the domestication syndrome’, iScience, 26(1). Available at: 10.1016/j.isci.2022.105704.

Vargas, J.Y., Fuenzalida, M. and Inestrosa, N.C. (2014) ‘In vivo activation of Wnt signaling pathway enhances cognitive function of adult mice and reverses cognitive deficits in an Alzheimer’s disease model’, Journal of Neuroscience, 34(6). Available at: 10.1523/JNEUROSCI.0862-13.2014.

Wang, Y.K. et al. (1999) ‘Characterization and expression pattern of the frizzled gene Fzd9, the mouse homolog of FZD9 which is deleted in Williams-Beuren syndrome’, Genomics, 57(2), pp. 235–248. Available at: 10.1006/geno.1999.5773.

Willfors, C. et al. (2021) ‘Williams syndrome: on the role of intellectual abilities in anxiety’, Orphanet Journal of Rare Diseases, 16(1). Available at: 10.1186/s13023-021-02098-4.

Williams Syndrome Association (2010) ‘Anxiety & Mood Problems in People with Williams Syndrome’. Available at: https://williams-syndrome.org/sites/williams-syndrome.org/files/Anxiety%26MoodProblemsinPeoplewithWS.pdf.

Xu, X. et al. (2020) ‘Wnt signaling in breast cancer: biological mechanisms, challenges and opportunities’, Molecular Cancer. Available at: 10.1186/s12943-020-01276-5.

Yde Ohki, C.M., et al. (2020) ‘The stress–Wnt-signaling axis: a hypothesis for attention-deficit hyperactivity disorder and therapy approaches’, Translational Psychiatry. Available at: 10.1038/s41398-020-00999-9.

Zanella, M. et al. (2019) ‘Dosage analysis of the 7q11.23 Williams region identifies BAZ1B as a major human gene patterning the modern human face and underlying self-domestication’, Science advances, 5(12), p. eaaw7908. Available at: 10.1126/sciadv.aaw7908.

Zhao, C. and Pleasure, S.J. (2005) ‘Frizzled9 protein is regionally expressed in the developing medial cortical wall and the cells derived from this region’, Developmental Brain Research, 157(1). Available at: 10.1016/j.devbrainres.2005.02.018.

Zhou, J. et al. (2022) ‘Atypical deletion of Williams–Beuren syndrome reveals the mechanism of neurodevelopmental disorders’, BMC Medical Genomics, 15(1), pp. 1–10. Available at: 10.1186/s12920-022-01227-7.

Zhou, W.J. et al. (2016) ‘The antidepressant roles of wnt2 and wnt3 in stress-induced depression-like behaviors’, Translational Psychiatry, 6(9). Available at: 10.1038/tp.2016.122.

Zwamborn, R.A.J. et al. (2018) ‘Wnt Signaling in the Hippocampus in Relation to Neurogenesis, Neuroplasticity, Stress and Epigenetics’, in Progress in Molecular Biology and Translational Science. Available at: 10.1016/bs.pmbts.2018.04.005.

