## Supplementary for "Disrupting *fzd9b* in zebrafish recapitulates stress- and anxiety-like behaviours associated with Williams syndrome"

SUPPLEMENTARY MATERIAL

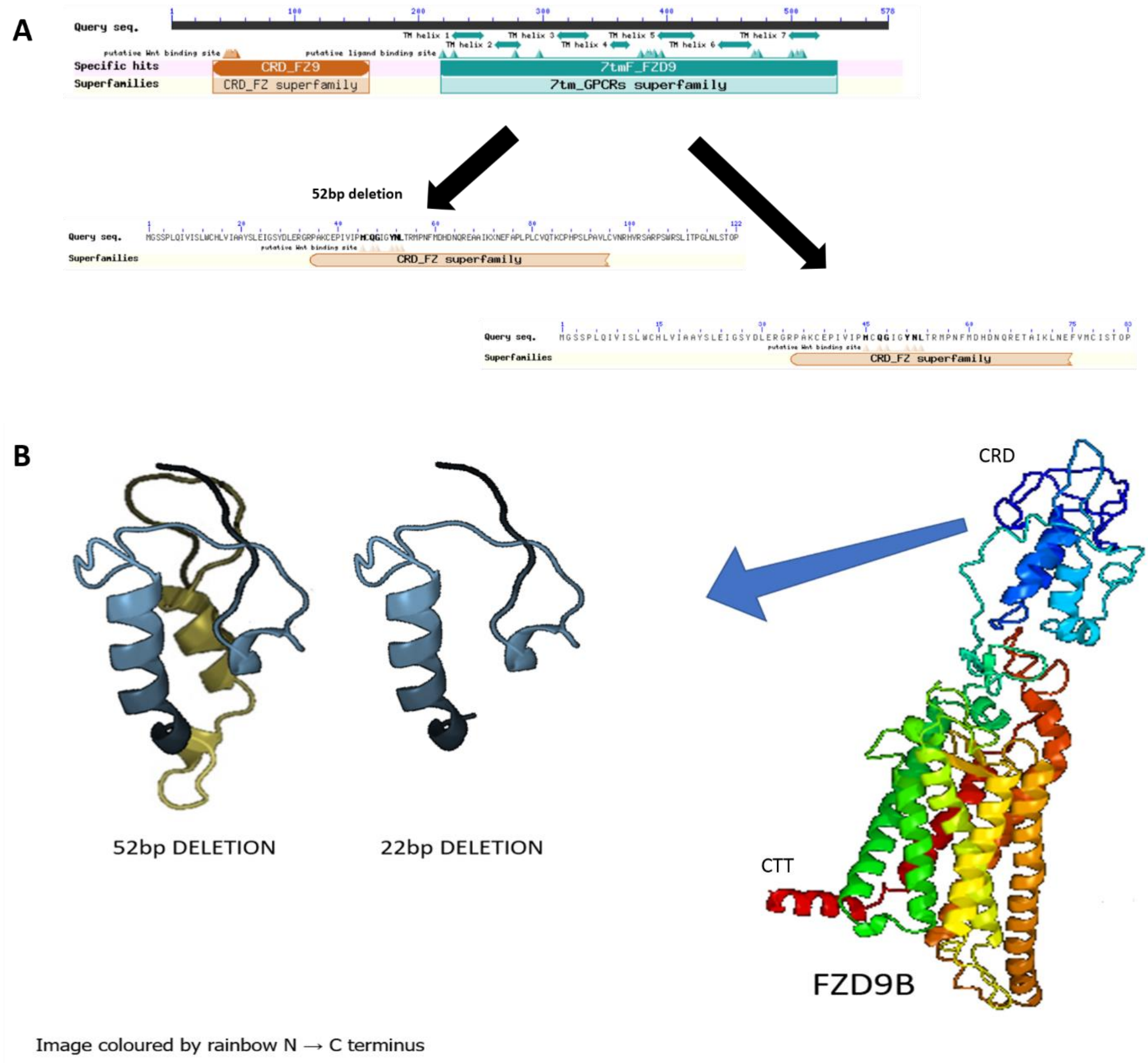


**Supplementary Figure 1. A)** Functional domains and **B)** 3D conformation for the Fzd9b receptor and the two small truncated proteins produced in the mutant lines. In both lines, the resulting truncated proteins only contain the Cysteine-rich Wnt-binding domain (CRD) while the intramembrane portion and the Carboxy-Terminal Tail (CTT) are absent. CRD is located extracellularly and is needed for binding Wnt proteins. Note that, although both truncated proteins are similar, the isolectric point (pI) of the 52 bp deletion is close to physiological levels (pI = 8) while the 22 bp differs (pI = 5). Furthermore, the additional 44 aa preceding the premature stop codon for the 52 bp deletion (dark yellow) codes for cysteines or unknown (X) aa. Functional domains were assessed with NCBI's CD-search tool (https://www.ncbi.nlm.nih.gov/Structure/cdd/wrpsb.cgi) and 3D structure with Phyre$^2$ (http://www.sbg.bio.ic.ac.uk/~phyre2/html/page.cgi?id=index).

**Supplementary Table 1.** Genes and corresponding primers used for qPCR.

| Gene name | ID number | forward (5'-3') | reverse (5'-3') | Length (bp) | Efficiency (%) | Source |
|---|---|---|---|---|---|---|
| *fzd9b* | ENSDARG00000014673 | TCCAGAATGGGAAGCTCACCTC | TCTCTCCAGGTCATAAGAGCCG | 99 | 100 | Own produced |
| *scn4aa* | ENSDART00000170961.2 | CGGCGATGTTTTGCATCAGTC | ACACGCAATTGGACAGGATGG | 114 | 99,1 | Own produced |
| *scn4ab* | ENSDART00000044157.8 | TGTTCATGGCGATGAGCGAC | GCAGAAACCTCTGGACAGAACC | 110 | 82 | Own produced |
| *tafa5l* | ENSDART00000133348.2 | TCAACCCAAGCGCACTATCG | CACACCACTGCTTGCTTTTCAC | 128 | 89 | Own produced |
| *wnt5a* | ENSDART00000164739.2 | AAAGACACTCCACACACACTGC | AGAGGATTCATCGCCAGAGACC | 107 | 88,6 | Own produced |
| *wnt5b* | ENSDART00000186829.1 | AGGACACCTACTTCTGGCAGTG | GGGGTTCATGGCTAATGACCAC | 91 | 97 | Own produced |
| *wnt9a* | ENSDART00000130349.3 | TTGAGAGATGGAACTGCACGC | GTGAGGCCCGCTGAAGAAATAG | 106 | 92,3 | Own produced |
| *ef1a* (housekeeping) | ENSDARG00000020850 | CTGGAGGCCAGCTCAAACAT | ATCAAGAAGAGTAGTACCGCTAGCATTAC | 87 | 88,4 | Doi:10.3389/fnbeh.2015,00271 |
| *rpll3α* (housekeeping) | ENSDARG00000044093 | TCTGGAGGACTGTAAGAGGTATGC | AGACGCACAATCTTGAGAGCAG | 148 | 80 | Doi:10.3389/fnbeh.2015,00271 |

**Supplementary Table 2.** Details of statistical analysis for the forced light dark transition assay performed in 5 days post fertilization larvae (WT, HET, HOM) for both *fzd9b*$^{22bpDEL}$ and *fzd9b*$^{52bpDEL}$ (Experiment 2).

| Line | Dependent variable | Period of the assay | Main effect | p value | χ2 | Degrees of freedom |
|---|---|---|---|---|---|---|
| ***fzd9b***$^{22bpDEL}$ | Distance travelled | Baseline | Time | p < 0.05 | 16.379 | 8 |
| ***fzd9b***$^{22bpDEL}$ | Distance travelled | Light 1 (min 10-20) | Time | p < 0.0001 | 32.4988 | 8 |
| ***fzd9b***$^{22bpDEL}$ | Distance travelled | Light 2 (min 30-40) | Time | p < 0.0001 | 97.694 | 8 |
| ***fzd9b***$^{22bpDEL}$ | Distance travelled | Light 3 (min 50-60) | Time | p < 0.0001 | 45.4471 | 8 |
| ***fzd9b***$^{22bpDEL}$ | Distance travelled | Dark 1 (min 20-30) | Time | p < 0.0001 | 81.2740 | 8 |
| ***fzd9b***$^{22bpDEL}$ | Distance travelled | Dark 2 (min 40-50) | Time | p < 0.0001 | 61.8538 | 8 |
| ***fzd9b***$^{22bpDEL}$ | Distance travelled | Dark 3 (min 60-70) | Time | p < 0.0001 | 63.6949 | 8 |
| ***fzd9b***$^{22bpDEL}$ | Slope of recovery | Light periods | Time | p < 0.05 | 554.916 | 29 |
| ***fzd9b***$^{52bpDEL}$ | Distance travelled | Baseline | Time | p < 0.0001 | 38.9689 | 8 |
| ***fzd9b***$^{52bpDEL}$ | Distance travelled | Light 1 (min 10-20) | Time | p < 0.0001 | 67.1889 | 8 |
| ***fzd9b***$^{52bpDEL}$ | Distance travelled | Light 2 (min 30-40) | Time | p < 0.0001 | 101.8495 | 8 |
| ***fzd9b***$^{52bpDEL}$ | Distance travelled | Light 3 (min 50-60) | Time | p < 0.0001 | 129..7344 | 8 |
| ***fzd9b***$^{52bpDEL}$ | Distance travelled | Dark 1 (min 20-30) | Time | p < 0.0001 | 115.1714 | 8 |
| ***fzd9b***$^{52bpDEL}$ | Distance travelled | Dark 2 (min 40-50) | Time | p < 0.0001 | 92.7437 | 8 |
| ***fzd9b***$^{52bpDEL}$ | Distance travelled | Dark 3 (min 60-70) | Time | p < 0.0001 | 114.3604 | 8 |
| ***fzd9b***$^{52bpDEL}$ | Slope of recovery | Light periods | Time | p < 0.05 | 114.3604 | 29 |

**Supplementary Table 3.** Details of statistical analysis for habituation and response to acoustic startle assay performed in 5 days post fertilization larvae (WT, HET, HOM) for both *fzd9b*$^{22bpDEL}$ and *fzd9b*$^{52bpDEL}$ in presence of the selective GSK-3 inhibitor BIO-acetoxime (200 µM). Baseline 1 refers to the basal locomotion before the flash of light, whereas baseline 2 refers to the basal locomotion after the flash of light and before the acoustic stimuli.

| **Line** | **Condition** | **Dependent variable** | **Period of the assay** | **Main effect** | **p value** | **χ2** | **Degrees of freedom** |
|---|---|---|---|---|---|---|---|
| ***fzd9b***$^{22bpDEL}$ | DMSO | Distance travelled | Baseline 1 | Time | $p < 0.05$ | 65.3706 | 59 |
| ***fzd9b***$^{22bpDEL}$ | DMSO | Distance travelled | Baseline 2 | Time | p < 2e-16 | 7354.1204 | 119 |
| ***fzd9b***$^{22bpDEL}$ | DMSO | Magnitude of response | Light flash | Time | p < 2e-16 | 115.773 | 1 |
| ***fzd9b***$^{22bpDEL}$ | DMSO | Slope of recovery | Light flash | Time | $p < 0.05$ | 6.1717 | 1 |
| ***fzd9b***$^{22bpDEL}$ | DMSO | Distance travelled | Taps | Genotype | $p < 0.05$ | 7.3771 | 2 |
| ***fzd9b***$^{22bpDEL}$ | DMSO | Distance travelled | Taps | Genotype * Time | p < 2e-16 | 148.0517 | 48 |
| ***fzd9b***$^{22bpDEL}$ | DMSO | Proportion of responders | Taps | - | $p > 0.05$ | - | - |
| ***fzd9b***$^{22bpDEL}$ | DMSO | Rate of habituation | Taps | Genotype | p < 3.972e-13 | 57.109 | 2 |
| ***fzd9b***$^{22bpDEL}$ | DMSO | Rate of habituation | Taps | Genotype * Stimulus number | p < 2.2e-16 | 554.916 | 27 |
| ***fzd9b***$^{22bpDEL}$ | BIO-acetoxime 200µM | Distance travelled | Baseline 1 | Time | $p < 0.05$ | 53.4153 | 59 |
| ***fzd9b***$^{22bpDEL}$ | BIO-acetoxime 200µM | Distance travelled | Baseline 2 | Time | p < 2e-16 | 29333.0044 | 119 |
| ***fzd9b***$^{22bpDEL}$ | BIO-acetoxime 200µM | Magnitude of response | Light flash | Time | p < 2e-16 | 127.4317 | 1 |

| | | | | | | | |
|---|---|---|---|---|---|---|---|
| ***fzd9b*** $^{22bpDEL}$ | BIO-acetoxime 200µM | Slope of recovery | Light flash | Time | $p < 0.001$ | 2.0533 | 1 |
| ***fzd9b*** $^{22bpDEL}$ | BIO-acetoxime 200µM | Distance travelled | Taps | Genotype * Drug * Time | $p < 0.05$ | 71.5187 | 48 |
| ***fzd9b*** $^{22bpDEL}$ | BIO-acetoxime 200µM | Proportion of responders | Taps | - | $p > 0.05$ | - | - |
| ***fzd9b*** $^{22bpDEL}$ | BIO-acetoxime 200µM | Rate of habituation | Taps | Genotype * Drug | $p < 7.653e-9$ | 37.847 | 4 |
| ***fzd9b*** $^{22bpDEL}$ | BIO-acetoxime 200µM | Rate of habituation | Taps | Genotype * Drug * Stimulus number | $p < 2.2e-16$ | 1076.527 | 29 |
| ***fzd9b*** $^{52bpDEL}$ | DMSO | Distance travelled | Baseline 1 | Time | $p < 0.01$ | 65.3706 | 59 |
| ***fzd9b*** $^{52bpDEL}$ | DMSO | Distance travelled | Baseline 2 | Time | $p < 2e-16$ | 7354.1204 | 119 |
| ***fzd9b*** $^{52bpDEL}$ | DMSO | Magnitude of response | Light flash | Time | $p < 2e-16$ | 115.773 | 1 |
| ***fzd9b*** $^{52bpDEL}$ | DMSO | Slope of recovery | Light flash | Time | $p < 0.05$ | 6.1717 | 1 |
| ***fzd9b*** $^{52bpDEL}$ | DMSO | Distance travelled | Taps | Time | $p < 2e-16$ | 686.671 | 24 |
| ***fzd9b*** $^{52bpDEL}$ | DMSO | Proportion of responders | Taps | - | $p < 0.05$ | - | - |
| ***fzd9b*** $^{52bpDEL}$ | DMSO | Rate of habituation | Taps | Genotype | $p < 2e-16$ | 338.3 | 2 |
| ***fzd9b*** $^{52bpDEL}$ | DMSO | Rate of habituation | Taps | Genotype * Time | $p < 2e-16$ | 1008.4 | 27 |
| ***fzd9b*** $^{52bpDEL}$ | BIO-acetoxime 200µM | Distance travelled | Baseline 1 | Time | $p < 0.05$ | 53.4153 | 59 |
| ***fzd9b*** $^{52bpDEL}$ | BIO-acetoxime 200µM | Distance travelled | Baseline 2 | Time | $p < 2e-16$ | 12819.9488 | 119 |

| | | | | | | | |
|---|---|---|---|---|---|---|---|
| ***fzd9b**$^{52bpDEL}$* | BIO-acetoxime 200µM | Magnitude of response | Light flash | Time | $p < 2e-16$ | 217.8463 | 1 |
| ***fzd9b**$^{52bpDEL}$* | BIO-acetoxime 200µM | Slope of recovery | Light flash | Time | $p < 0.05$ | 7.4217 | 1 |
| ***fzd9b**$^{52bpDEL}$* | BIO-acetoxime 200µM | Distance travelled | Taps | Time | $p < 2e-16$ | 1332.3594 | 24 |
| ***fzd9b**$^{52bpDEL}$* | BIO-acetoxime 200µM | Proportion of responders | Taps | - | $p > 0.05$ | - | - |
| ***fzd9b**$^{52bpDEL}$* | BIO-acetoxime 200µM | Rate of habituation | Taps | Drug | $p < 9.555e-9$ | 37.414 | 4 |
| ***fzd9b**$^{52bpDEL}$* | BIO-acetoxime 200µM | Rate of habituation | Taps | Genotype * Drug * Stimulus number | $p < 2e-16$ | 893.929 | 29 |
